# Extended Pavlovian conditioned approach training and devaluation in Long-Evans rats using time-restricted feeding

**DOI:** 10.64898/2026.11.04.764351

**Authors:** Ivanka L. Rainer, Kyle Brooks, Yvonne M. Ulrich-Lai, Jayme R. McReynolds

## Abstract

Pavlovian conditioned approach (PCA) is a useful tool in studying compulsive- and addictive-like behaviors. PCA pairs a cue with reward delivery and rats typically differentiate into three phenotypes (sign-trackers, goal-trackers, and intermediates) based on behavior during the reward-paired cue presentation. Sign-trackers, who preferentially interact with the cue, are more likely to engage in compulsive behaviors and have decreased behavioral flexibility compared to goal-trackers, who preferentially interact with the food cup. However, one limitation to PCA is the use of food deprivation conditions that maintain rats at 80-90% of *ad libitum* weight, likely increasing stress which may bias animals towards sign-tracking behavior. Furthermore, most behavioral analysis only examines lever pressing and food cup entry, overlooking other behaviors that may better define phenotypes. In the current study male and female Long-Evans rats underwent an extended PCA training paradigm followed by satiety-induced reward devaluation under time-restricted feeding (TRF) conditions. Under TRF, rats continued to gain weight across PCA training and maintained caloric intake relative to *ad libitum* intake. Expanding the scope of behavioral analysis increased the proportion of ST rats in both early and late PCA training. Rats also increased the time spent interacting with the food cup during the inter-trial interval across extended PCA training. All rats were sensitive to satiety-induced reward devaluation, regardless of sex or devaluation condition. This study helps establish a PCA method optimized for Long-Evans rats with relevance for identifying PCA phenotypes contribution to dysregulated behavior in the context of chronic stress and obesity.

## 1.1 Introduction

A key part of survival is learning what environmental cues predict reward or threat, thereby identifying opportunities to seek food and avoid risk. However, this association of cues with specific outcomes may also become maladaptive. In cases of obsessive-compulsive disorder (OCD), for example, conditioned cues act as triggers for compulsive behaviors (Marzuki et al., 2024). In gambling addiction, these associations lead to suboptimal and impulsive decision making and reinforce maladaptive reward-seeking behavior (Swintosky et al., 2022). Environmental cues may also cause an aversive appetitive response, triggering drug-seeking or food-seeking in the absence of hunger (Flagel et al., 2009; Horstmann et al., 2015).

Pavlovian conditioned approach (PCA), or Pavlovian autoshaping, has been a useful tool in studying compulsive- and addiction-like behaviors. PCA involves repeated pairing of a cue consisting of lever extension and light presentation followed by the response-independent delivery of a reward (e.g., a sucrose pellet). During repeated PCA testing, animals quickly learn to discriminate between the conditioned stimulus reward-paired trials (CS+) vs the unpaired trials (CS-). Rodents typically segregate into three behavioral phenotypes based on their behavior during the CS+ cue presentation: sign-trackers (ST) – interacting mostly with the cue; goal-trackers (GT) –interacting mostly with the location of reward delivery (ie. the food cup); and intermediate (INT) – equally likely to interact with both cue and reward locations (Fitzpatrick et al., 2013; Meyer et al., 2012). Importantly, these PCA phenotypes have important implications for other reward-related behaviors. Sign trackers are more likely to over-engage in rewarding-seeking behaviors and have a higher vulnerability for substance use disorders (Kuhn et al., 2018; Swintosky et al., 2022). Furthermore, sign trackers display behavioral inflexibility and are less sensitive to reward devaluation (either through satiety or aversion) than goal trackers (Bien & Smith, 2023; Keefer et al., 2020; Morrison et al., 2015; Stapf et al., 2024). While individual rats seem to have a natural predisposition toward one of these three phenotypes, longer durations of PCA training tends to shift all rats towards more sign-tracking behavior (Bien & Smith, 2023; Flagel et al., 2009; Keefer et al., 2020; Stapf et al., 2024).

However, there are limitations to the current model of PCA, as PCA is typically performed on animals undergoing marked food deprivation to maintain them at 80-90% of *ad libitum* body weight (Bacharach et al., 2023; Bien & Smith, 2023; Derman & Matthew Lattal, 2023; Stapf et al., 2024). This food restriction is used to increase motivation for the food reward used during the task. However, given that stress increases the likelihood of sign-tracking behavior, it is possible that the stress associated with strict food deprivation could bias animals towards sign tracking behavior (Derman & Matthew Lattal, 2023; Lomanowska et al., 2011; Turfe et al., 2024). Additionally, the food deprivation conditions limits examination of PCA behavior under different dietary conditions such as high fat diet exposure. Furthermore, PCA behavior is often quantified through automated measurements of lever presses and head entries into a food cup (Bien & Smith, 2023; Derman & Matthew Lattal, 2023; Meyer et al., 2012; Stapf et al., 2024; Turfe et al., 2024). However, this overlooks the possible presence of other behaviors that may be occurring during cue presentation but do not trigger a lever press or head entry – such as sniffing or staring intensely at the lever or cup or investigating the cue light – and does not consider the behavior that occurs during the inter-trial interval. The present study aims to resolve the limitations through an extended PCA protocol on male and female Long Evans rats maintained on a less severe, time-restricted feeding schedule that allowed free access to food during a specific 4-h time window each day. Additionally, the present study expands the behavioral analysis to examine total interaction with the cue or food cup in contrast to the traditional use of lever presses/food cup entry for determination of PCA scores.

## 1.2 Methods

### 1.2.1 Animals

Adult Long-Evans rats, females approximately 200-225g and males approximately 250-275g, were purchased from Envigo (Indianapolis, IN). Rats were single housed in a temperature and humidity-controlled housing facility with a 12/12 reverse light cycle (lights off at 09:00h, lights on at 21:00h). Housing facilities were accredited by the Association for the Assessment and Accreditation of Laboratory Animal Care (AAALAC). Animals acclimated for at least one week in the animal facilities before the experiment started. All procedures were approved by the University of Cincinnati Institutional Animal Care and Use Committee and conducted in compliance with NIH guidelines.

### 1.2.2 Food access

Rats received *ad libitum* access to a normal chow diet (Teklad LM-485, Envigo, Madison, WI, USA; 17% daily calories from fat, 25% from protein, and 58% from carbohydrate with no added sucrose). Body weight and food intake was measured for 3 days prior to the onset of time-restricted feeding. For time-restricted feeding access to food, rats received daily 4-hour access (13:00h-17:00h) to normal chow. This time window was chosen because it allowed at least 1hr between the end of the previous PCA session, to avoid association with the completion of pavlovian conditioned approach (PCA) training and the return of food, and at least 16hr of fasting prior to the following PCA session to motivate behavior during PCA training. Time-restricted feeding began 10 days before the start of PCA training (Fig. 1) to allow rats time to acclimate to the feeding schedule and body weight and food intake was recorded daily at the time of food removal. Food intake was recorded as total grams as well as kcal. Normalized food intake was also calculated as kcal/day/body weight.

**Figure 1.**
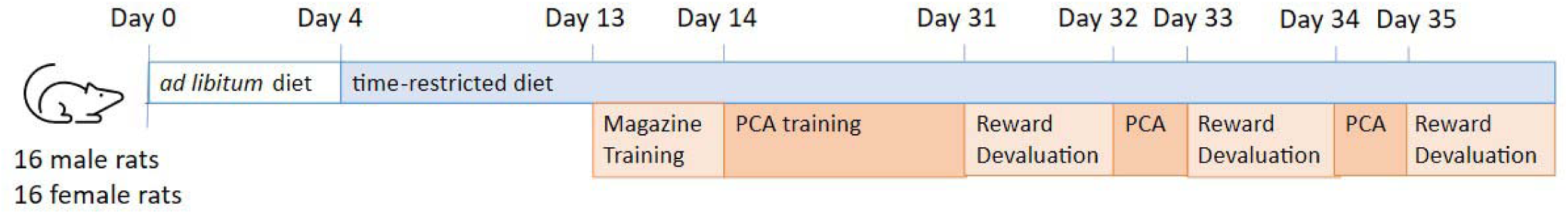
Schematic representation of the experiment timeline. Rats were maintained on ad lib normal chow diet for days 1-3. Time-restricted feeding (4 h/day) began on day 4 and continued through the rest of the experiment. Magazine training took place on day 13. Starting on day 14, the Pavlovian conditioned approach (PCA) task began and continued for 17 days. Rats then underwent reward devaluation testing on days 31, 33, and 35, with a day of normal PCA testing in between each devaluation day. Three different reward devaluation conditions were tested in each rat (non-devalued, food satiety, sensory-specific satiety) and these were presented in a counter-balanced order.

### 1.2.3 Pavlovian Conditioned Approach

All experiments were completed using Med Associated operant conditioning chambers encased in sound attenuating cubicles with a ventilation fan (MED-Associates Inc., St Albans, VT). Each chamber was equipped with two retractable levers on one wall of the chamber, a cue light above each lever, an external food cup situated between the two levers, a house light on the back wall, a food hopper for delivery of sucrose pellets, a Pi USB Cam (Hou & Glover, 2022) above the chamber, and two infrared lights situated on the back wall of the sound attenuating cubicle to allow for video analysis of approach behaviors.

Rats were pre-exposed to several 45 mg unflavored sucrose pellets (45mg, Bio-Serv, Flemington, NJ; 3.84 kcal/gram) in their home cage at least three times during the week prior to PCA training to limit neophobia. The amount of sucrose pellets consumed was recorded to include in the assessment of daily food intake. One day prior to the onset of PCA training, rats underwent magazine training (between 9:30-11:00h) to acclimate rats to sucrose delivery and consumption from the food-cup. Rats were each placed into the operant chamber and received a total of 25 sucrose pellets delivered over a randomized interval of 15-45s. Any sucrose remaining after the session was recorded.

The following day, rats began once-daily 30-min PCA training sessions (between 09:30-12:00h; adapted from *Spring et al.* (2021)), consisting of 50 total trials. For 25 trials, a lever was extended and the cue light above the lever was illuminated for 10s after which a response-independent sucrose pellet was delivered (conditioned stimulus (CS)+ trial). For the other 25 trials, the opposite lever was extended and the cue light above the lever was illuminated for 10s but was not followed by sucrose delivery (CS-trial). Trials were presented in a random order and separated by a variable intertrial interval averaging 30 sec (range of 15-45s). Lever presses were recorded but sucrose delivery was non-contingent. Additionally, the session was recorded with Pi USB cameras for later behavioral analysis. At the conclusion of each training session, the food cup was checked to confirm that all delivered sucrose pellets were eaten. Rats underwent PCA training for 17 days.

Behavior during PCA training was recorded and manually scored using BORIS (Behavioral Observation Research Interactive Software; Friard & Gamba, 2016). As the rats engaged in a significant amount of lever-directed behavior beyond lever presses, these behaviors were scored and quantified. Lever-directed behavior was defined as sniffing, staring, biting, grabbing, or investigating the lever or cue light. Food cup-directed behavior was defined as sniffing, staring, biting, grabbing, or investigating the food cup. The number of interactions with the lever/light or the food cup, duration of the interactions, latency to approach either the lever/light or the food cup were quantified for each trial and ethograms of the behaviors across each PCA session were produced. Behavior during the inter-trial interval was assessed for days 5, 10 and 17 of PCA training. Lever presses were recorded and extracted from Med-PC using MedPC to excel software. One metric of learning used for the PCA task was the probability of approach, calculated as (the number of trials of a given type (CS+ or CS-) in which at least one lever/light or food cup interaction was made/number of trials of the corresponding type). The probability of approach was calculated for each day of PCA training. Another metric of learning for the PCA task was the discrimination index, calculated as the difference between approach probability to either the cue or food cup during the CS+ cue vs. the CS-cue normalized to overall probability: (P_CS+_ – P_CS-_)/(P_CS+_ + P_CS-_). A positive discrimination index indicates a preference for approaching the CS+, a negative discrimination index indicates a preference for the CS-, and a discrimination index of zero indicates equal likelihood of approach during either the CS+ or CS-trials.

Rats were determined to be sign-trackers, goal-trackers, or intermediates using a PCA score (Meyer et al., 2012), which averages three behavioral indices: (1) response bias, which compares time spent in lever-vs. food cup-directed behavior (time spent with lever – time spent with food cup)/(time spent with lever + time spent with food cup); (2) latency bias, which compares the average latency from cue presentation to onset of lever-vs. food cup-directed behavior (lever latency – food cup latency)/10; and (3) probability difference, which compares the probability of lever-vs. food cup-directed behavior during a cue presentation (P_lever_ – P_food cup_). As defined by *Meyer et al.* (2012), sign-trackers were categorized as having a PCA score >0.5, goal-trackers as having a PCA score <-0.5, and intermediates as having a PCA score between 0.5 and -0.5. PCA scores were calculated for each day of PCA training and early PCA training was defined as day 5 and late PCA training was defined as day 17 of PCA training. Traditionally response bias is calculated using lever presses or food cup head entry rather than total time interacting with the lever/light or the food cup. The differences in PCA score distribution and assignment of behavioral phenotype using the time spent interacting with the lever/light or food cup (all interaction analysis) and lever press/food cup head entry were compared at day 5 and day 17.

### 1.2.4 Reward Devaluation

After the last day of PCA training, rats underwent reward devaluation to determine how behavioral flexibility differed between males and females after extended PCA training. To further examine sensory-specific satiety, each rat was tested under 3 different devaluation conditions: non-devalued, chow-induced satiety, and sucrose-induced satiety, with a day of PCA training in between each devaluation day. The order of devaluation received was counterbalanced between rats. Devalued rats were given either a large quantity of sucrose or access to a food hopper one hour prior to training to induce satiety. Rats then underwent a PCA session under extinction (no sucrose pellets were dispensed following CS+ trials). For reward devaluation behavior the total time interacting with either the lever/light or the food cup, the probability of approach to either the lever/light or the food cup, and the total number of interactions with the lever/light or the food cup were assessed. Following the reward devaluation session, all rats were given access to sucrose pellets for 15 minutes and the amount of sucrose consumed was recorded to confirm satiety.

### 1.2.5 Statistical Analysis

Data are shown as mean ± SEM. Statistical analyses were conducted using GraphPad Prism (San Diego, CA). When comparing multiple variables/groups, data were analyzed with two-or three-way ANOVA, with repeated measures when appropriate (RM-ANOVA). When comparing two groups, unpaired t-tests were used. All ANOVAs were followed by Holm-Šídák or Dunnett’s post hoc testing where appropriate. Statistical significance was set as p<0.05. All significant main and interactive effects are reported in the results section with accompanying F-or t-values and p-values.

## 1.3 Results

### 1.3.1 Time-restricted feeding for 4 h/day did not decrease body weight or caloric intake

Rats were given *ad libitum* access to normal chow diet for four days to establish baseline body weight and caloric intake. Rats were then placed on a time-restricted feeding schedule where they received daily 4-hour access to normal chow diet. A two-way repeated measures (RM) ANOVA analysis of body weight across experimental days revealed a significant time x sex interaction (Fig. 2A; F (33,990) = 35.88, p<0.0001), as well as significant main effects of sex (F (1,30) = 347.20, p<0.0001) and time (F (33,990) = 91.73, p<0.0001). A Dunnett’s post hoc test showed that male rats showed increased body weight, as compared to Day 1 body weight, starting on day 7 and continuing throughout PCA training (p<0.05). In females, a Dunnett’s post hoc test showed increased body weight, as compared to Day 1 body weight, starting on day 13 and continuing throughout PCA training (p<0.05). Therefore, male rats began showing increases in body weight during the initial time-restricted feeding period and continued to show increases in body weight throughout the PCA training period. Female rats had much more modest increases in body weight and did not exhibit significant increases in body weight until the end of the initial time-restricted feeding period but maintained these increases throughout the PCA training period. A two-way RM ANOVA analysis of caloric intake (kcal) across experimental days revealed a significant time x sex interaction (Fig. 2B; F (32,960) = 3.50, p<0.0001) and significant main effects of sex (F (1,30) = 152.70, p<0.0001) and time (F (32,960) = 18.69, p<0.0001). A Dunnett’s post hoc test indicated that male rats showed some significant decreases in caloric intake during the initial time-restricted feeding period (days 4, 6, 7, 10, 13; p<0.01) as compared to day 1 intake but then maintained caloric intake throughout the PCA training period. A Dunnett’s post hoc test indicated that female rats did not show significant changes in caloric intake throughout the initial time-restricted feeding period as compared to day 1 intake and actually showed modest significant increases in caloric intake at various points throughout the PCA period (days 18, 20, 22, 25, 28; p<0.05). As male rats were overall larger than female rats, daily caloric intake was also assessed as normalized to body weight (kcal/day/BW; Fig. 2C). A two-way RM ANOVA revealed a significant time x sex interaction (F (32,960) = 3.00, p<0.0001) and a significant main effect of time (F (32, 960) = 15.43, p<0.0001). A Holm-Šídák post hoc test showed that male rats showed greater normalized caloric intake prior to the time-restricted feeding paradigm than females (days1-3, p<0.01). Taken together, these data indicate that the time-restricted feeding paradigm allows for rats to be in a food deprived state immediately prior PCA daily while maintaining normal daily caloric intake and body weight gain.

**Figure 2.**
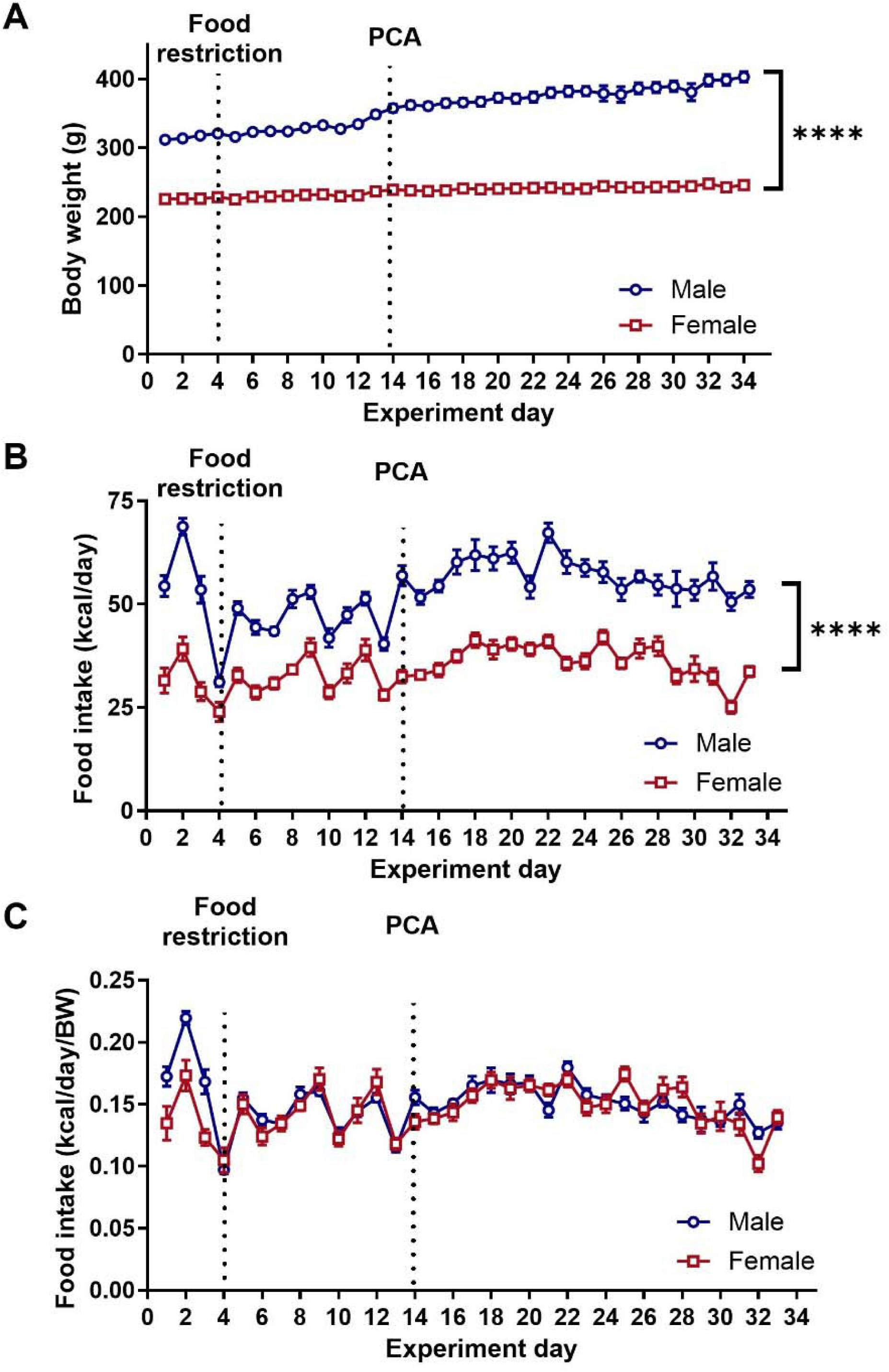
Body weight and caloric intake were relatively unchanged by 4-hr time-restricted feeding in both male and female rats. Time course of (A) body weight, (B) caloric intake, and (C) caloric intake normalized to body weight across experimental days. Vertical dotted lines at days 4 and 14 indicate the onset of the time-restricted feeding and PCA paradigms, respectively. (A) Male and female rats increased their body weight across the time-restricted feeding period with males having higher overall weight and greater weight gain across the experimental period. (B) At the onset of the time-restricted feeding paradigm caloric intake (kcal/day) dropped briefly but quickly returned to *ad libitum* levels throughout PCA training and male rats consumed more food than female rats. (C) When food intake was normalized to body weight, male and female rats consumed similar quantities of food each day though male rats showed greater normalized caloric intake prior to the onset of the time-restricted feeding paradigm. Additionally, the time-restricted feeding paradigm did not significantly impact overall food intake compared to *ad libitum* levels. Data are shown as mean ± SEM. n=16 rats/group. ****p<0.0001 indicates a main effect of sex. Additionally, there was a significant time x sex interaction (p<.0001 for body weight and caloric intake).

### 1.3.2 Acquisition of PCA

All rats quickly learned to discriminate between the CS+ and CS-cues. The number of CS+ lever presses was greater than the number of CS-lever presses starting on the third day of the PCA paradigm (Fig. 3A). A three-way RM ANOVA did not reveal a significant day x CS type x sex interaction (F (16,960) = 1.49, p=0.10), but did reveal a significant CS type x sex interaction (F (1,60) = 7.51, p<0.01), a significant day x CS type interaction (F (16,960) = 6.24, p<0.0001), and significant main effects of CS type (F (1,60) = 73.46, p<0.001), sex (F (1,60) = 7.11, p<0.01), and day (F (16,960) = 3.23, p<0.0001), with male rats having more lever presses overall in response to the CS+ compared to females once the PCA paradigm had been learned. A Holm-Šídák post hoc test indicated that the number of CS+ lever presses was significantly greater than the CS-lever presses starting on the third day of the PCA paradigm, indicating successful discrimination between the two cues. Similarly, rats had a higher probability of approach during the CS+ than the CS-cue starting on the second day of the PCA paradigm (Fig. 3B). A three-way RM ANOVA did not reveal a significant day x CS type x sex interaction (F (16,960) = 0.93, p=0.54) but did reveal a significant day x CS type interaction (F (16,960) = 12.66, p<0.0001), and significant main effects of day (F (16,960) = 4.41, p<0.001), CS type (F (1,60) = 93.12, p<0.0001), and sex (F (1,60) = 6.54, p<0.05). Male rats had a greater probability of approach in response to the CS+ cue overall than females after the PCA paradigm had been learned. A Holm-Šídák post hoc test revealed that the probability of approach in response to the CS+ cue was significantly greater compared to the CS-cue starting on day 2 of the PCA training period. Male and female rats showed differences in overall lever presses in response to the CS+ cue; therefore, to correct for these differences in overall responding at the lever, a discrimination index was calculated to compare the ratio of probability of approach to *either* the cue or food cup in response to the CS+ cue vs. the CS-cue (Fig. 3C). A two-way RM ANOVA did not reveal a significant day x sex interaction (F (16,480) = 0.76, p=0.73) or a significant main effect of sex (F (1,30) = 1.98, p=0.17), but did reveal a significant main effect of day (F (16,480) = 7.78, p<0.0001), indicating that the effects of sex on the number of lever presses was primarily due to a difference in the overall degree of lever responding and not indicative of differences in the acquisition of the PCA task.

**Figure 3.**
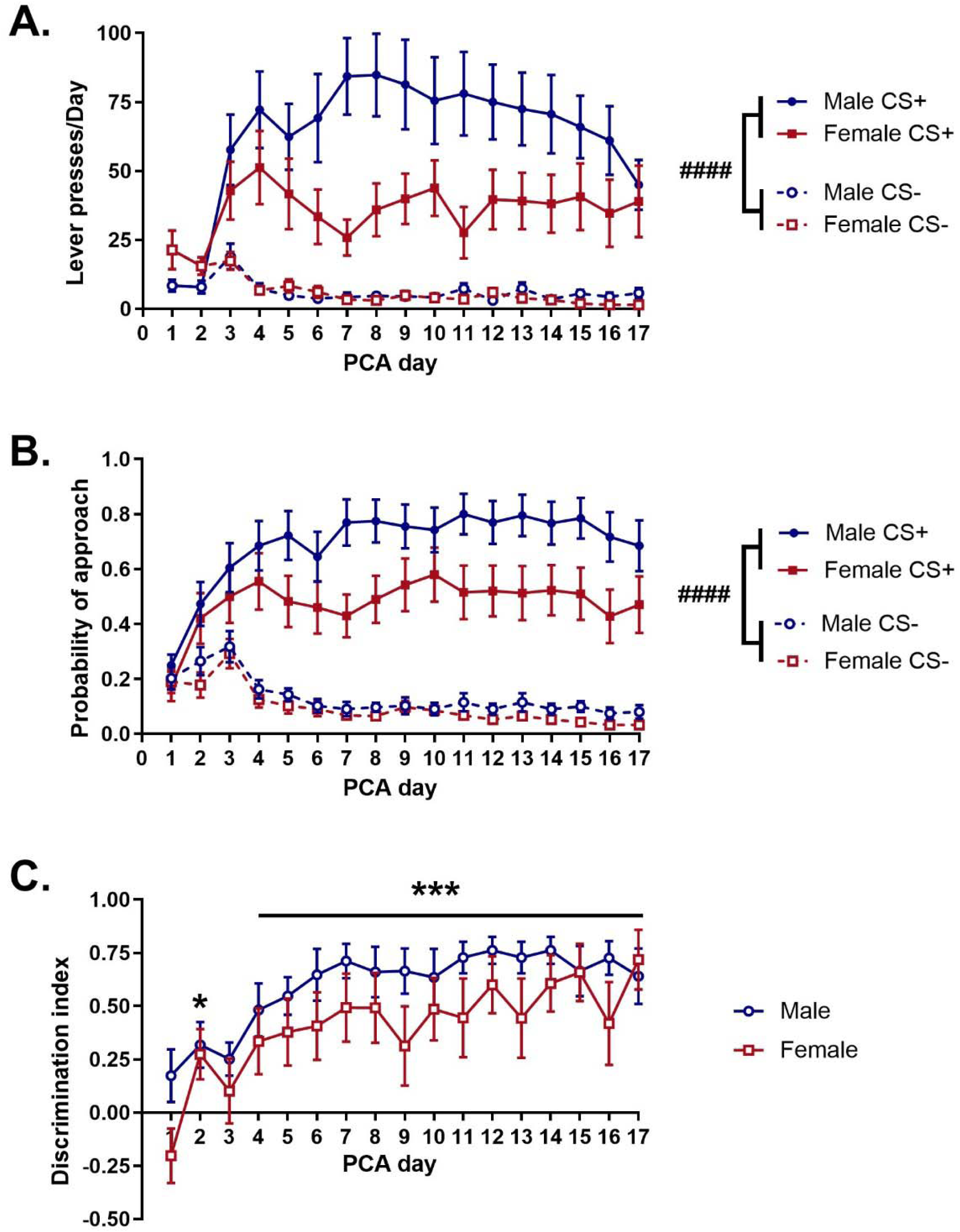
Male and female rats quickly learned to discriminate between CS+ cue and CS-cue during the Pavlovian conditioned approach (PCA) task while male rats exhibited greater overall responding to the CS+ cue than female rats. A) While male and female rats showed differentiation in lever presses in response to the CS+ and CS-cue by day 3 (males) – 4 (females), male rats had greater overall lever responding to the CS+ cue than female rats throughout the PCA task. B) Male and female rats quickly showed significant differences in the probability of approach to the CS+ or the CS-cue by day 2 of the PCA. However, males showed a greater overall probability of approach to the CS+ cue than female rats. C) Male and female rats showed similar increases in the discrimination index (difference between approach probability to either the cue or food cup during the CS+ or CS-trial normalized to overall probability) across training days. There was a significant increase in the discrimination index on day 2 and days 4-17 compared to day 1. Data are shown as mean ± SEM. n=16 rats/group. ####p<0.0001; *p<0.05, ***p<0.001, compared to day 1. There is a significant main effect of sex for both lever presses/day (p<0.01) and probability of approach (p<0.05).

### 1.3.3 Behavioral phenotypes shifted toward increased sign-tracking representation with extended PCA training and the method of analysis changed the distribution of PCA scores in early and late PCA training

Behavior was scored across the 17 days of the PCA paradigm. PCA scores were calculated using total interaction time with the lever/light or food cup (all interaction analysis) for the response bias. As PCA training progresses, PCA scores increase in both males and females, indicating a general shift towards sign tracking behavior (Fig. 4A). The method of analysis for calculating PCA scores impacts the overall distribution of the behavioral phenotype. We observed that Long Evans rats engaged in multiple lever/light-or food cup-directed behaviors beyond lever presses or food cup entry such as sniffing, staring, biting, grabbing, or investigating the lever/light or food cup. Therefore, we compared the distribution of PCA scores using the traditional method involving lever presses/food cup entry only or using the total interaction time with the light/lever or food cup (all interaction analysis). In early PCA training (day 5) of PCA training, an unpaired two-tailed t-test revealed no significant differences in PCA score between males and females using either the all interaction analysis approach (Fig. 4B left; t(30) = 0.11, p=0.91) or the lever press/food cup entry analysis approach (Fig. 4B right; t(30) = 1.17, p=0.25). However, the method of analysis did alter the distribution of behavioral phenotypes with a Fisher’s exact test revealing a trend towards a significant change in the distribution of PCA scores (p=0.08). In males, the all interaction analysis approach resulted in 1 goal trackers (GT), 6 Intermediates (INT), and 9 sign trackers (ST). However, the lever press/food cup entry analysis approach resulted in 1 GT, 11 INT, and 4 ST. In females, the all interaction analysis approach resulted in 1 GT, 4 INT, and 11 ST and the lever press/food cup entry analysis approach resulted in 0 GT, 5 INT, and 11 ST. The method of analysis did not alter the distribution of behavioral phenotypes in female rats. A two-way RM ANOVA comparing PCA scores based on analysis approach in both male and female rats in early PCA training (day 5) revealed a significant analysis approach x sex interaction (Fig. S1A; F (1,30) = 4.752, p<0.05) and a significant main effect of analysis approach (F (1,30) = 6.57, p<0.05). A Fischer’s post hoc test revealed significantly higher PCA scores using the all interaction analysis compared to the lever press/food cup entry analysis in male rats only (p<0.01). In early PCA training (day 5), the all interaction analysis approach captured more sign tracking behavior, yielding a greater proportion of sign trackers, in male rats only.

**Figure 4.**
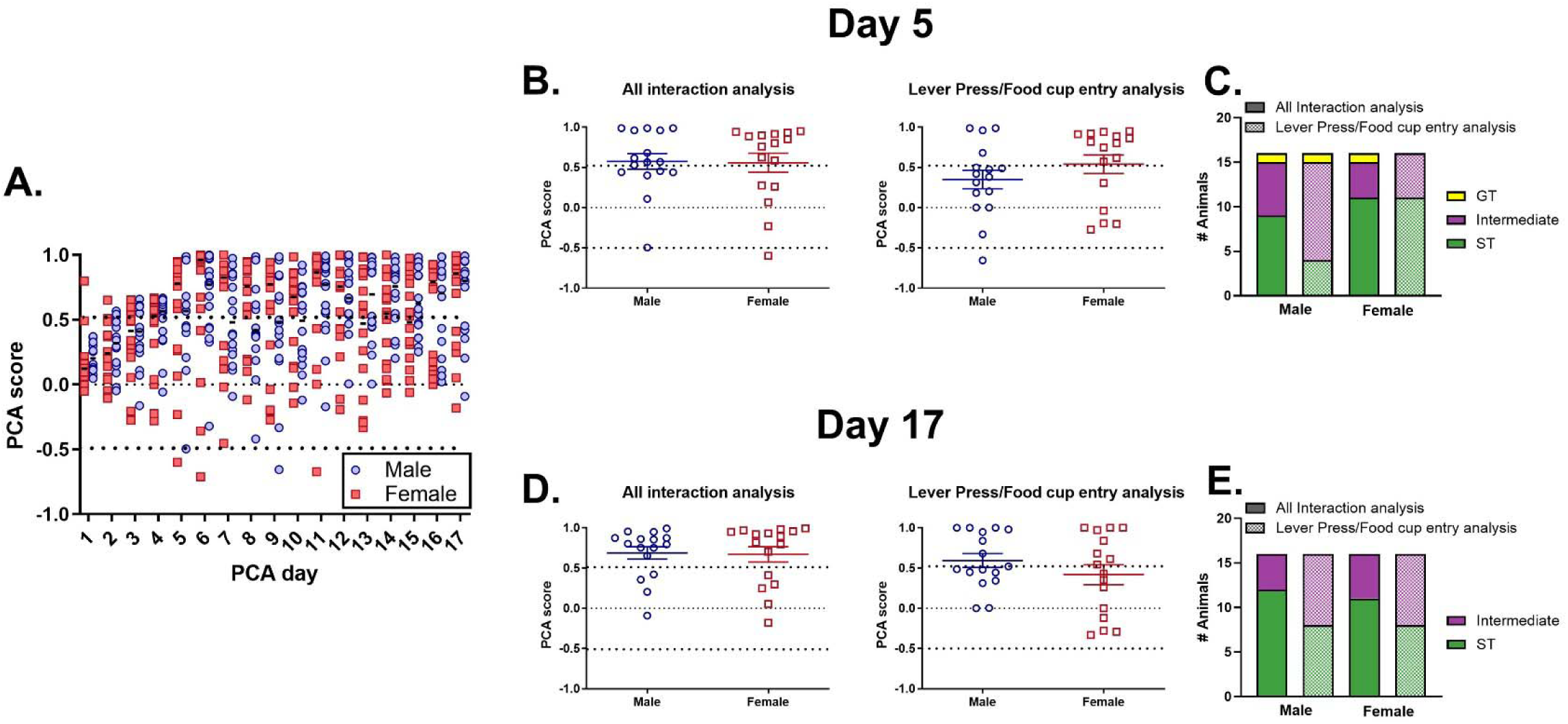
Pavlovian conditioned approach (PCA) score increased with extended PCA training regardless of sex, indicating a shift towards greater sign-tracking behavior but the distribution of behavioral phenotypes was impacted by analysis method. (A) Scatterplot of PCA scores for individual rats across days of PCA training in both male and female rats using an analysis that considers all interactions with the CS+ cue or food cup beyond lever presses and food cup entries. B) Distribution of PCA scores on day 5 of PCA training in male and female rats using either all interaction analysis (left) or lever press/food cup entry analysis (right). The distribution of PCA scores was not significantly different between male and female rats using either analysis method. C). The method of analysis did change the overall distribution of behavioral phenotypes on day 5 with the all interaction analysis producing a greater number of sign-trackers in males (Fisher’s exact test, p=0.054). All interaction analysis: Goal trackers (GT): Male – 0; Female – 1; Intermediate (INT): Male – 6; Female – 4; Sign trackers (ST): Male – 10; Female – 11. Lever press/food cup entry analysis: GT: Male -1; Female – 0; INT: Male – 11; Female – 5; ST: Male – 4; Female – 11. D) Distribution of PCA scores on day 17 of PCA training in male and female rats using either all interaction analysis or lever press/food cup entry analysis. The distribution of PCA scores was not significantly different between males and females using either analysis method. E) The analysis method significantly changed the distribution of behavioral phenotypes in both male and female rats (Fisher’s exact test, p<0.0001). Similar to day 5, all interaction analysis produced a greater number of sign trackers than lever press/food cup analysis. However, in contrast to day 5 this was observed in both male and female rats. All interaction analysis: GT: Male – 0; Female – 0; INT: Male – 4; Female – 5; ST: Male – 12; Female – 11. Lever press/food cup entry analysis: GT: Male – 0; Female – 0; INT: Male – 8; Female – 8; ST: Male – 8; Female – 8. n=16 rats/group.

In late PCA training (day 17), an unpaired two-tailed t-test revealed no significant differences in PCA scores between males and females using either the all interaction analysis approach (Fig. 4D left; t(30) = 0.14, p=0.89) or the lever press/food cup entry analysis approach (Fig. 4D right; t(30) = 1.71, p=0.25). However, a Fisher’s exact test revealed that method of analysis did significantly shift the distribution of behavioral phenotypes. In males, the all interaction analysis approach resulted in 0 GT, 4 INT, and 12 ST whereas the lever press/food cup entry analysis approach resulted in 0 GT, 8 INT, and 8 ST. In females, the all interaction analysis approach resulted in 0 GT, 5 INT, and 11 ST whereas the lever press/food cup entry analysis approach resulted in 0 GT, 8 INT, and 8 ST. A two-way RM ANOVA comparing PCA scores based on analysis approach in both male and female rats in late PCA training (day 17) did not reveal a significant analysis approach x sex interaction (Fig. S1B; F (1,30) = 2.63, p=0.12) but did reveal a significant main effect of analysis approach (F (1,30) = 13.83, p<0.0001). Therefore, in both males and females, the all interaction analysis approach captured more sign tracking behavior resulting in a higher proportion of sign trackers compared to the lever press/food cup entry analysis approach in late PCA training.

Given the observed differences in PCA score distribution by analysis approach, we examined the individual behavioral indices that comprise the PCA score (Fig. S2; response bias, latency bias, and probability difference). Response bias was examined using either the all interaction analysis approach or the lever press/food cup entry analysis approach. In early PCA training (day 5), unpaired two-tailed t-tests did not reveal significant differences between male and female rats in response bias using all interaction analysis (Fig. S2A; t(30)=0.30, p=0.77), response bias using lever press/food cup entry analysis (Fig. S2B; t(30) = 1.20, p=0.24), latency bias (Fig. S2C; t(30) = 0.52, p=0.61), or probability difference (Fig. S2D; t(30) = 1.26, p=0.22). A two-way RM ANOVA comparing response bias in male and female rats based on analysis approach (Fig. S3A) did not reveal a significant analysis approach x sex interaction (F (1,30) = 0.76, p=0.39), a significant main effect of analysis approach (F (1,30) = 0.26, p=0.12), or a significant main effect of sex (F (1,30) = 1.13, p=0.30). However, while there was no significant effect of analysis type on response bias scores, it is likely that changes in this metric did drive the change in distribution of PCA scores observed in male rats in early PCA training (Fig. 4C). We also examined these same behavioral indices in late PCA training (day 17). An unpaired two-tailed t-test did not reveal significant differences between male and female rats in response bias using all interaction analysis (Fig. S2E; t(30)=0.49, p=0.63), response bias using lever press/food cup entry analysis (Fig. S2F; t(30) = 1.44, p=0.16), latency bias (Fig. S2G; t(30) = 0.57, p=0.57), or probability difference (Fig. S2H; t(30) = 0.55, p=0.59).). A two-way RM ANOVA comparing response bias in male and female rats based on analysis approach (Fig. S3B) did not reveal a significant analysis approach x sex interaction (F (1,30) = 0.76, p=0.39) or a significant main effect of sex (F (1,30) = 1.40, p=0.25) but did reveal a significant main effect of analysis approach (F (1,30) = 13.83, p=0.0001). Therefore, the changes in the distribution of behavioral phenotypes in male and female rats in late PCA training (day 17) because of analysis approach are explained by the shifts in response bias calculations. These differences in response bias calculations lead to higher proportions of sign tracker rats in late PCA training using the all interaction analysis approach.

### 1.3.4 Rats showed multiple types of cue-directed behaviors during CS+ cue presentation in early and late PCA training

Typically, during the PCA task, cue-directed behavior is quantified as lever presses, while cup-directed behavior is quantified as head entries into the food-cup (Bien & Smith, 2023; Derman & Matthew Lattal, 2023; Meyer et al., 2012; Stapf et al., 2024; Turfe et al., 2024). However, we observed many other cue-directed behaviors (e.g., biting, grabbing, or sniffing of the lever or light) that did not trigger a lever press, suggesting that the conventional index of cue-directed behavior may not fully capture the full extent of cue-directed behaviors. As such, we compared cue-directed behaviors across PCA phenotypes (sign-trackers, goal-trackers, and intermediate rats) when defined by lever presses alone vs. an expanded behavioral analysis that included all cue-directed behaviors (Fig. 5). However, we did not include goal trackers in the analysis due to the very low n (n=2) in early PCA training. In early PCA training, ST rats lever press during the CS+ trials more than INT rats and female rats generally lever press less than males (Fig. 5A). A two-way ANOVA did not reveal a significant phenotype (ST vs INT) by sex interaction on lever presses (F (1,26) = 0.04, p=0.84) but did reveal a significant main effect of phenotype (F (1,26) = 7.70, p<0.05), and a significant main effect of phenotype (F (1, 26) = 11.07, p<0.01). While ST rats display more lever pressing than INT rats, INT rats generally have a larger number of lever/light interactions than ST rats, but this effect is primarily driven by the female rats (Fig. 5B). A two-way ANOVA did reveal a significant phenotype x sex interaction on the number of lever/light interactions (F (1,26) = 9.54, p<0.01) and significant main effects of phenotype (F (1,26) = 4.86, p<0.05), and sex (F (1,26) = 33.33, p<0.0001). A Holm-Šídák post hoc test indicated that female INT rats have a larger number of lever/light interactions than female ST rats (p<0.0001), but that difference is not observed in male rats (p>0.05). Additionally, female INT rats have a greater number of lever/light interactions than male INT rats (p<0.05). ST rats spend more time overall interacting with the lever/light than INT rats (Fig. 5C). A two-way ANOVA did not reveal a significant phenotype x sex interaction (F (1,26) = 2.82, p=0.11) or a significant main effect of sex (F (1,26) = 0.89, p=0.35) but did reveal a significant main effect of phenotype (F (1,26) = 68.47, p<0.0001). ST rats also had a greater average bout length for lever/light interactions than INT rats (Fig. 5D). A two-way ANOVA did reveal a significant phenotype x sex interaction (F (1,26) = 5.99, p<0.05) and a significant main effect of phenotype (F (1,26) = 84.31, p<0.0001) but not a significant main effect of sex (F (1,26) = 0.89, p=0.35). A Holm-Šídák post hoc test indicated that female and male ST rats have longer average lever/light interactions bouts than INT rats (p<0.001 for each sex). Additionally, there was a trend for female INT rats to have longer lever/light interaction bouts than male INT rats (p=0.10). Taken together, ST rats spend more time interacting with the lever/light through fewer but longer bouts of interaction whereas INT rats interact with the lever/light more frequently but for shorter periods of time, particularly in female rats.

**Figure 5.**
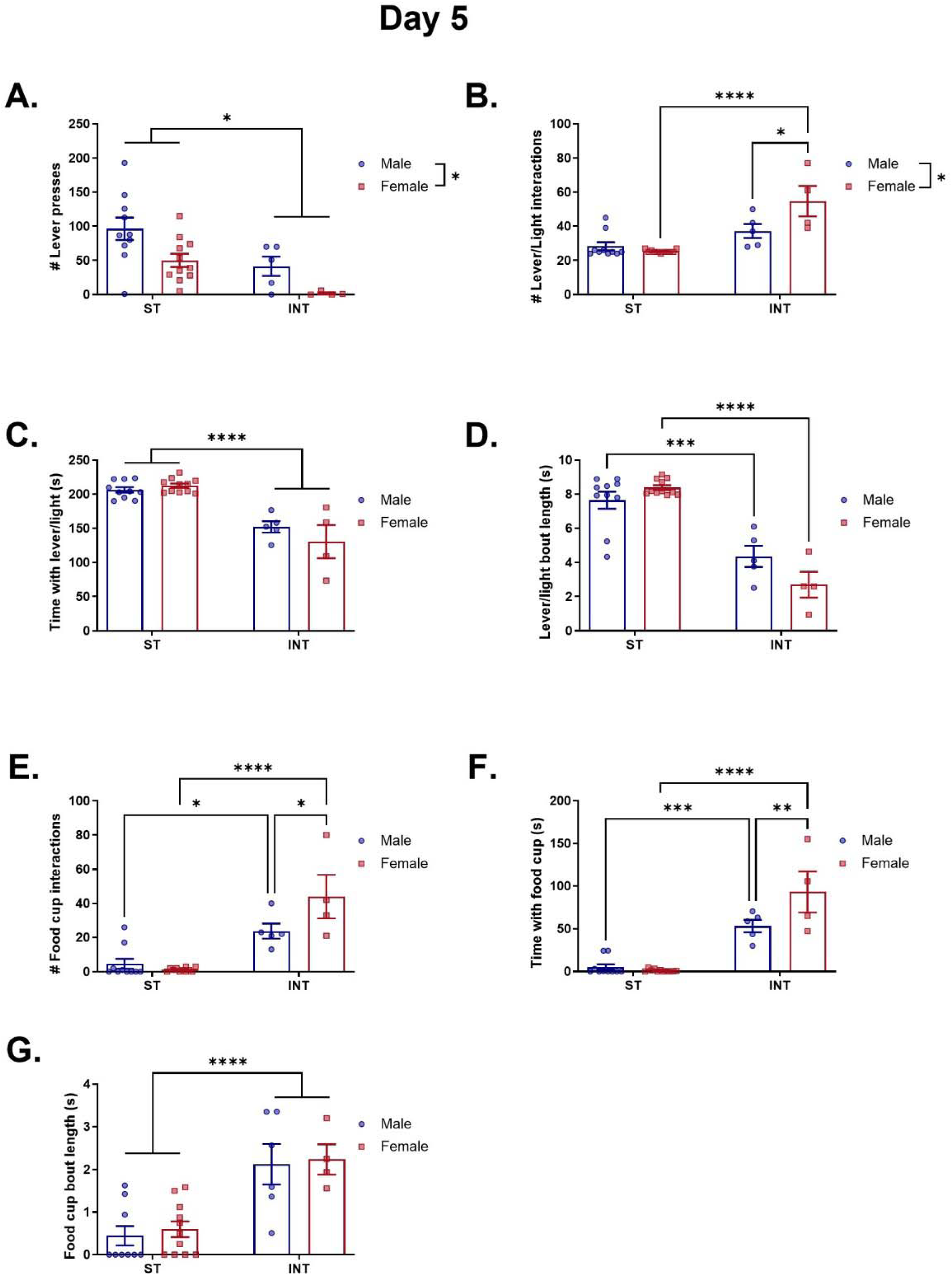
Sex- and behavioral phenotype-specific responses during the CS+ trial on day 5 of PCA training. A) While sign tracker rats (ST) showed greater lever presses than intermediate (INT) rats, males showed greater overall lever presses during the CS+ trial than female rats. B). INT rats showed greater overall number of interactions with the CS+ cue (lever and light) beyond lever presses (sniffing, staring, biting, grabbing, or investigating the lever or cue light) than ST rats. Female INT rats had a significantly greater number of lever/light interactions than male rats. C) ST male and female rats spent a greater amount of time overall interaction with the lever/cue light than INT rats. D) ST male and female rats had a longer lever/light interaction bout length compared to INT rats. E) INT rats had a greater number of food cup interactions (sniffing, staring, biting, grabbing, or investigating the food cup) than ST rats. Female INT rats has a greater number of food cup interactions compared to male INT rats. F) INT rats spent more time interacting with the food cup compared to ST rats with female INT spending more time interacting with the food cup than male INT rats. G). INT rats had a longer food cup interaction bout length compared to ST rats with male and female rats demonstrating similar bout lengths in either phenotype. Data are shown as mean ± SEM. *p<0.05, **p<0.01, ***p<0.001, ****p<0.0001.

In early PCA training (day 5), INT rats have a greater number of interactions with the food cup than ST rats with female INT rats showing the highest amount of food cup interactions (Fig. 5E). A two-way ANOVA revealed a significant phenotype x sex interaction (F (1,26) = 7.27, p<0.05) and a significant main effect of phenotype (F (1,26) = 50.13, p<0.0001) but not a significant main effect of sex (F (1,26) = 3.67, p=0.07). A Holm-Šídák post hoc test indicated that both female and male ST rats have a greater number of food cup interactions than INT rats (male: p<0.05; female: p<0.0001). Additionally, female INT rats have a greater number of food cup interactions than male INT rats (p<0.05). INT rats spend more time overall with interacting with the food cup than ST rats with female INT rats spending the greatest amount of time with the food cup (Fig. 5F). A two-way ANOVA revealed a significant phenotype x sex interaction (F (1,26) = 8.86, p<0.01) and a significant main effect of phenotype (F (1,26) = 89.62, p<0.0001) and sex (F (1,26) = 5.90, p<0.05). A Holm-Šídák post hoc test indicated that both female and male ST rats spend more time interacting with the food cup than INT rats (p<0.001 for both sexes). Additionally, female INT rats spend more time interacting with the food cup than male INT rats (p<0.01). INT rats have longer average bouts of interactions with the food cup than ST rats in both male and female rats (Fig. 5G). A two-way ANOVA did not reveal a significant phenotype x sex interaction (F (1,26) = 0.01, p=0.95) or a significant main effect of sex (F (1,26) = 0.19, p=0.67) but did reveal a significant main effect of phenotype (F (1,26) = 29.48, p<0.0001). Taken together, INT rats interact more frequently and for longer than ST rats, as expected, though female INT rats have the greatest number of interactions and spend the most time with the food cup compared to male INT rats and ST rats. In early PCA training, ST rats spend the majority of their time at the lever/light as indicated by their higher lever pressing, lower number of interactions but longer interaction time and spend little to no time at the food cup during each CS+ trial (Fig. 6A). While the data was not graphed for GT rats, they spend most of their time at the food cup and for longer periods of time though they do still engage with the lever/light during some CS+ trials (Fig. 6C). Interestingly, INT rats seem to demonstrate more of a switching behavior between the lever/light and the food cup during CS+ trials (Fig. 6B). They have a greater number of interactions with the lever/light than ST rats but the length of time they spend at the lever/light is lower.

**Figure 6.**
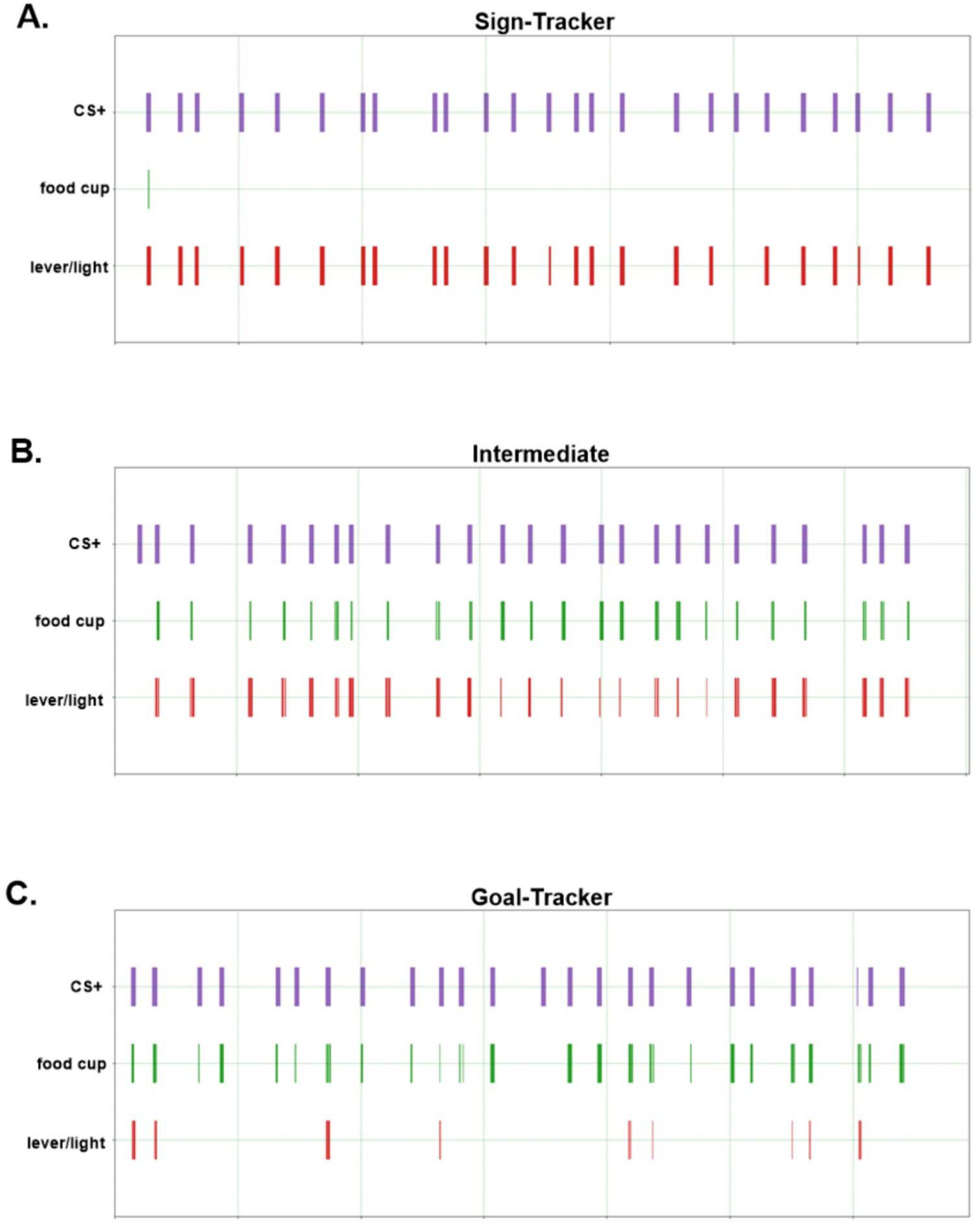
Representative ethograms illustrate the pattern of lever- and cup-directed behaviors during each CS+ presentation across the behavioral phenotypes on day 5 of PCA training. An ethogram is shown for a representative (A) sign-tracker, (B) intermediate, and (C) goal-tracker rat. The top row of each ethogram indicates the timing/duration of each CS+ presentation, while the middle and bottom rows indicate food cup and cue/lever directed behaviors, respectively. Sign tracker rats direct almost all of their directed behavior towards the lever/light and goal tracker rats direct the majority of their directed behavior towards the food cup. However, intermediate rats switch between the food cup and the lever/light.

In late PCA training (day 17), there was no significant difference in lever pressing during the CS+ trials across phenotype or sex unlike that observed in early PCA training (Fig. 7A). A two-way ANOVA did not reveal a significant phenotype (ST vs INT) by sex interaction on lever presses (F (1,28) = 2.29, p=0.14) or significant main effects of phenotype (F (1,28) = 1.18, p=0.29) or sex (F (1, 28) = 0.89, p=0.35). INT rats have a greater number of lever/light interactions than ST rats in late PCA training but unlike early PCA training male and female INT behavior is similar (Fig. 7B). A two-way ANOVA did not reveal a significant phenotype x sex interaction on the number of lever/light interactions (F (1,28) = 1.39, p=0.25) or a significant main effect of sex (F (1,28) = 1.07, p=0.31) but did reveal a significant main effect of phenotype (F (1,26) = 5.88, p<0.05). Like early PCA training, ST rats spend more time overall interacting with the lever/light than INT rats in late PCA training (Fig. 7C). A two-way ANOVA did not reveal a significant phenotype x sex interaction (F (1,28) = 0.72, p=0.40) or a significant main effect of sex (F (1,28) = 0.81, p=0.38) but did reveal a significant main effect of phenotype (F (1,28) = 188.50, p<0.0001). Like early PCA training, ST rats also had a greater average bout length for lever/light interactions than INT rats in late PCA training (Fig. 7D). A two-way ANOVA did not reveal a significant phenotype x sex interaction (F (1,28) = 0.21, p=0.65) or a significant main effect of sex (F (1,28) = 0.30, p=0.59) but did reveal a significant main effect of phenotype (F (1,28) = 57.05, p<0.0001). Taken together, ST rats spend more time interacting with the lever/light through fewer but longer bouts of interaction whereas INT rats interact with the lever/light more frequently but for shorter periods of time. This pattern of behavior is similar to what is observed in early PCA training though in late PCA training female INT rats now perform similarly to male INT rats unlike early PCA training.

**Figure 7.**
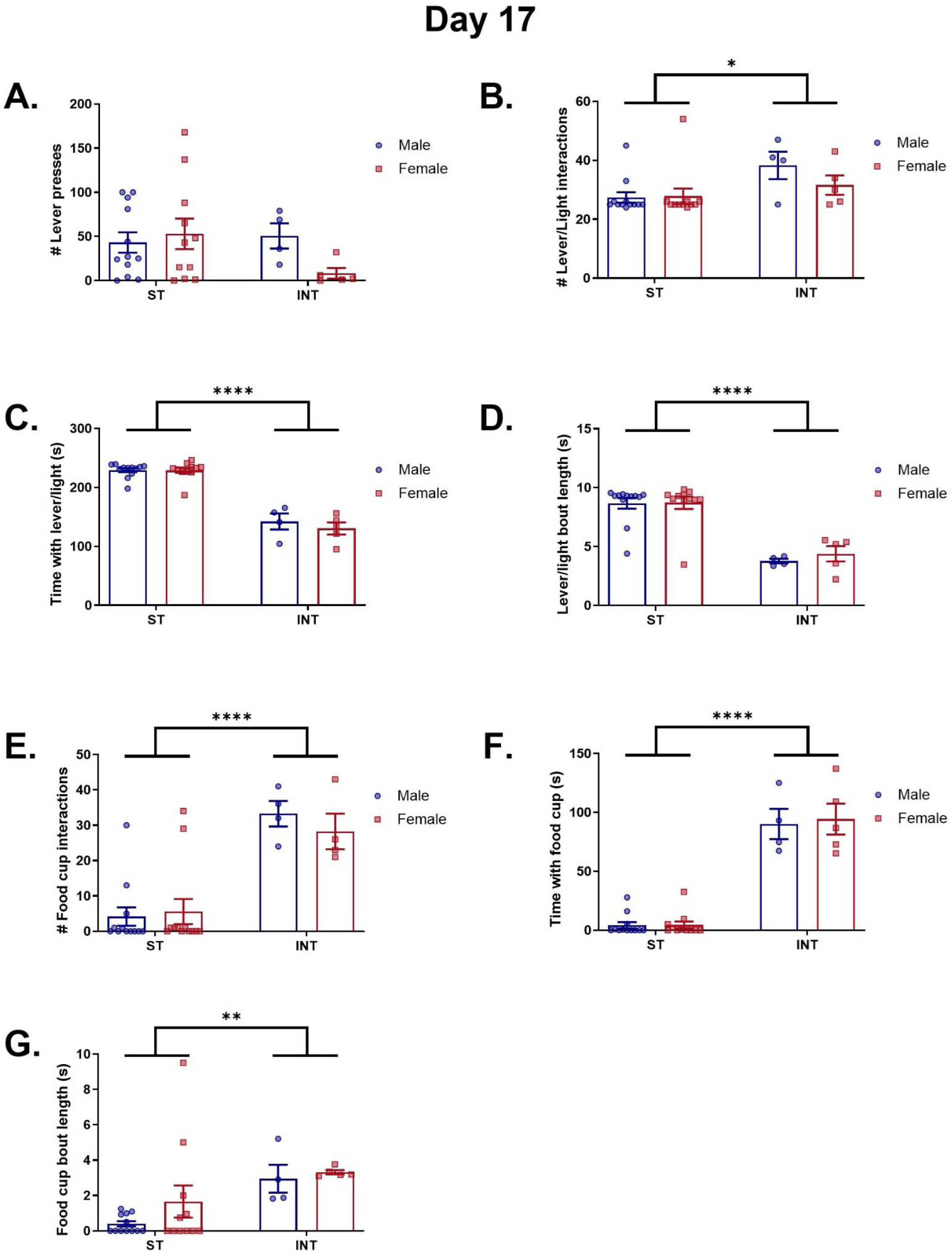
Sex- and behavioral phenotype-specific responses during the CS+ trial on day 17 of PCA training. A) There were no sex or behavior phenotype differences on the number of lever presses during the CS+ trials on day 17 of PCA training. B). INT rats showed greater overall number of interactions with the CS+ cue (lever and light) beyond lever presses (sniffing, staring, biting, grabbing, or investigating the lever or cue light) than sign tracker (ST) rats. C) ST male and female rats spent a greater amount of time overall interaction with the lever/cue light than INT rats. D) ST male and female rats had a longer lever/light interaction bout length compared to INT rats. E) INT rats had a greater number of food cup interactions (sniffing, staring, biting, grabbing, or investigating the food cup) than ST rats. F) INT rats spent more time interacting with the food cup compared to ST rats. G). INT rats had a longer food cup interaction bout length compared to ST rats with male and female rats demonstrating similar bout lengths in either phenotype. Data are shown as mean ± SEM. **p<0.01, ****p<0.0001.

In late PCA training (day 17), like early PCA training, INT rats have a greater number of interactions with the food cup than ST rats (Fig. 7E). A two-way ANOVA did not reveal a significant phenotype x sex interaction (F (1,28) = 0.58, p=0.45) or a significant main effect of sex (F (1,28) = 0.18, p=0.67) but did reveal a significant main effect of phenotype (F (1,28) = 37.85, p<0.0001). Like early PCA training, INT rats spend more time overall with interacting with the food cup than ST rats in late PCA training (Fig. 7F). A two-way ANOVA did not reveal a significant phenotype x sex interaction (F (1,28) = 0.09, p=0.76) or a significant main effect of sex (F (1,28) = 0.12, p=0.73) but did reveal a significant main effect of phenotype (F (1,28) = 191.8, p<0.0001). Like early PCA training, INT rats have longer average bouts of interactions with the food cup than ST rats in both male and female rats in late PCA training (Fig. 7G). A two-way ANOVA did not reveal a significant phenotype x sex interaction (F (1,28) = 0.36, p=0.56) or a significant main effect of sex (F (1,28) = 1.16, p=0.29) but did reveal a significant main effect of phenotype (F (1,28) = 7.87, p<0.01). Taken together, INT rats interact more frequently and for longer than ST rats, as expected, in late PCA training. However, unlike early PCA training, female INT rats interact with the food cup with a similar frequency and overall time compared to male INT rats.

### 1.3.5 Rats increase their time spent interacting with the food cup during the inter-trial interval across extended PCA training, particularly in female rats

Behavior during the inter-trial interval (ITI) was also examined on days 5, 10, and 17 of the extended PCA paradigm. The time spent interacting with the CS+ location, time spent interacting with the food cup, and time spent interacting with the CS-location was quantified. The time spent interacting with the CS+ location during the inter-trial interval was unchanged with extended training (Fig. 8A). A two-way RM ANOVA did not reveal a significant day x sex interaction (F (2,60) = 1.78, p=0.77), or main effects of day (F (2,60) = 0.55, p=0.58) or sex (F (1,30) = 0.48, p=0.49). However, time spent interacting with the food cup increased with extended training and was overall greater in female rats (Fig. 8B). A two-way RM ANOVA did not reveal a significant day x sex interaction (F (2,60) = 0.93, p=0.40) but did reveal significant main effects of day (F (2,60) = 8.48, p<0.001) and sex (F (1,30) = 10.71, p<0.05). A Holm-Šídák post hoc that time spent interacting with the food cup was greater on day 17 compared to day 5 (p<0.001) and day 10 (p<0.01). The time spent interacting with the CS-location during the inter-trial interval was unchanged with extended PCA training (Fig. 8C). A two-way RM ANOVA did not reveal a significant day x sex interaction (F (2,60) = 0.11, p=0.89), or main effects of day (F (2,60) = 1.26, p=0.29) or sex (F (1,30) = 1.04, p=0.32). The rats spent most of the time during the inter-trial interval interaction with the food cup which increased with additional days of PCA training and female rats generally spent more time with the food cup during the ITI than male rats.

**Figure 8.**
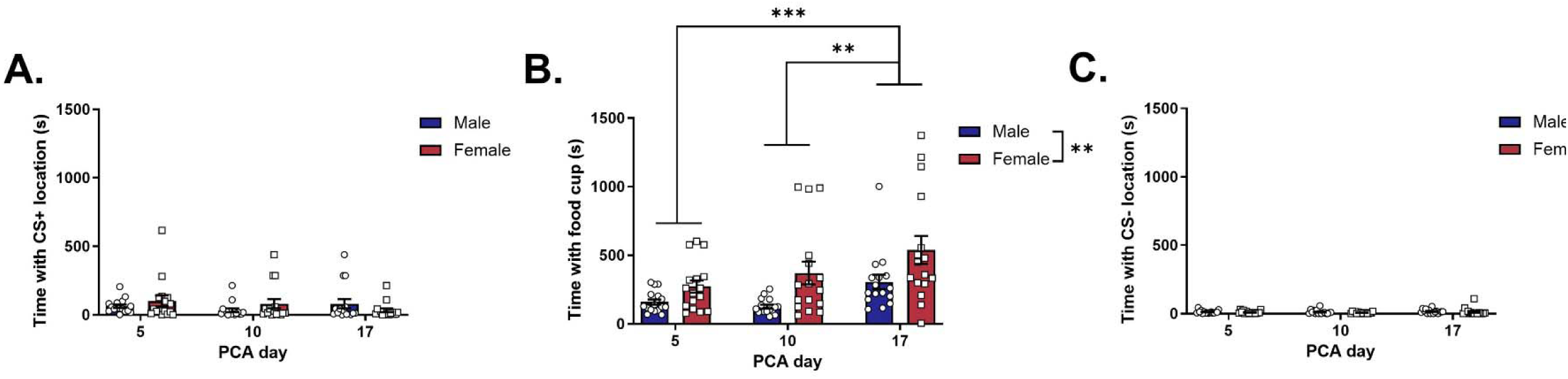
Time spent interacting with the food-cup during the inter-trial interval (ITI) increased with extended PCA training, particularly in female intermediate rats. A) The time spent with the CS+ location during the ITI between trials does not differ from early to late PCA training or by sex. B) Rats spend most of their time during the ITI interacting with the food cup and the time spent with the food cup increases from early to late PCA training. Overall female rats spend more time interacting with the food cup during the ITI than male rats. C). Male and female rats spend very little time with the CS-location during the ITI between trials. This time does not change from early to late PCA training nor does it differ by sex. Data are shown as mean ± SEM. **p<0.01, ***p<0.001.

**Figure 9.**
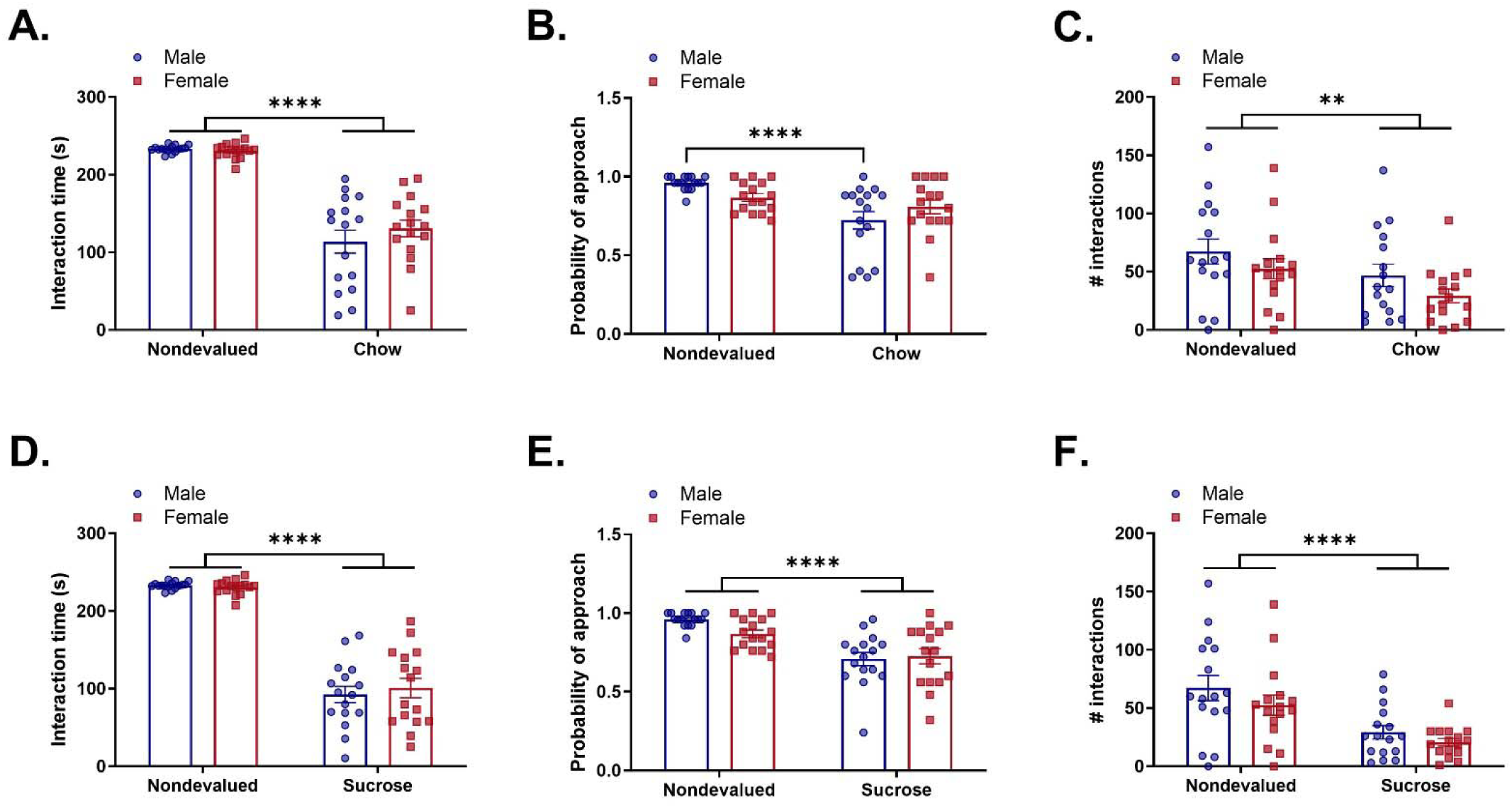
PCA responding was sensitive to reward devaluation by both chow and sucrose pre-exposure. A) Time spent interacting with either the lever/light or the food cup was decreased by satiety induced by pre-testing access to standard chow. The degree of devaluation was similar between male and female rats. B) Unlike total interaction time, only male rats showed a significant decrease in the probability of approach to either the food cup or lever/light following chow-induced satiety. Female rats did not show a significant decrease in probability of approach with chow-induced satiety. C) Both male and female rats showed a decrease in the number of interactions with either the lever/light or the food cup following chow-induced satiety. D). Time spent interacting with either the lever/light or the food cup was decreased by sensory-specific satiety with sucrose. The degree of devaluation was similar between male and female rats. E) Sensory-specific satiety with sucrose decreased the probability of approach to either the food cup or the lever/light in both male and female rats. F) Sensory-specific satiety with sucrose decreased the number of interactions with either the lever/light or the food cup in both male and female rats. Data are shown as mean ± SEM. **p<0.01, ****p<0.0001.

### 1.3.6 Reward devaluation

To test sensitivity to reward devaluation, rats underwent PCA testing under 3 different devaluation conditions: (1) devalued with standard chow; (2) devalued with sucrose pellets; and (3) no devaluation. Behavior during the devaluation test was assessed by examining interaction time, probability of approach, and the number of interactions with either the lever/light or the food cup. Satiety-induced devaluation with pre-exposure to chow decreased the interaction time with the lever/light and food cup in both male and female rats. A two-way RM ANOVA did not reveal a significant devaluation condition (nondevalued vs chow) x sex interaction (F (1,30) = 1.26, p=0.27) or a significant main effect of sex (F (1,30) = 0.56, p=0.46) but did reveal a significant main effect of devaluation condition (F (1,30) = 157.7, p<0.0001). Satiety-induced devaluation with pre-exposure to chow decreased the probability of approach in male but not female rats. A two-way RM ANOVA revealed a significant devaluation condition x sex interaction (F (1,30) = 5.65, p<0.05) and a significant main effect of devaluation condition (F (1,30) = 15.86, p<0.001) but not a significant main effect of sex (F (1,30) = 0.01, p=0.92). A Fisher’s post hoc test indicated that probability of approach was significantly reduced in the chow devaluation condition compared to the nondevalued condition in male (p<0.001) but not female (p>0.05) rats. Satiety-induced devaluation with pre-exposure to chow decreased the total number of interactions with the lever/light and food cup in both male and female rats. A two-way RM ANOVA did not reveal a significant devaluation condition (nondevalued vs chow) x sex interaction (F (1,30) = 0.04, p=0.85) or a significant main effect of sex (F (1,30) = 2.35, p=0.14) but did reveal a significant main effect of devaluation condition (F (1,30) = 9.71, p<0.01).

Sensory-specific satiety induced by pre-exposure to sucrose decreased the interaction time with the lever/light and food cup in both male and female rats. A two-way RM ANOVA did not reveal a significant devaluation condition (nondevalued vs chow) x sex interaction (F (1,30) = 0.46, p=0.50) or a significant main effect of sex (F (1,30) = 0.11, p=0.74) but did reveal a significant main effect of devaluation condition (F (1,30) = 288.9, p<0.0001). Sensory-specific satiety-induced devaluation with pre-exposure to sucrose decreased the probability of approach to the lever/light and food cup in both male and female rats. A two-way RM ANOVA did not reveal a significant devaluation condition (nondevalued vs chow) x sex interaction (F (1,30) = 2.46, p=0.13) or a significant main effect of sex (F (1,30) = 1.18, p=0.29) but did reveal a significant main effect of devaluation condition (F (1,30) = 31.65, p<0.0001). Sensory-specific satiety-induced devaluation with pre-exposure to sucrose decreased the total number of interactions with the lever/light and food cup in both male and female rats. A two-way RM ANOVA did not reveal a significant devaluation condition (nondevalued vs chow) x sex interaction (F (1,30) = 0.32, p=0.58) or a significant main effect of sex (F (1,30) = 1.56, p=0.22) but did reveal a significant main effect of devaluation condition (F (1,30) = 39.63, p<0.0001). Pre-exposure to chow or sucrose successfully produced reward devaluation across all behavioral indices following extended PCA training. However, female rats in the chow-induced devaluation condition did not show significant decreases in probability of approach but did in the sucrose-induced devaluation condition.

## 1.4 Discussion

The present work explored the effects of extended Pavlovian conditioned approach (PCA) training on behavioral phenotype classification and reward devaluation in male and female Long Evans rats on a time-restricted feeding schedule. While most studies use food restriction to a defined limited amount of food daily to keep rats at approximately 80-90% of their original starting weight, the 4-hr feeding window with time-restricted feeding allowed rats to maintain *ad libitum* body weight and continue weight gain throughout PCA training. Importantly, rats effectively learned the PCA task and were able to differentiate between the CS+ and CS-within 3 days of training indicating that the greater caloric consumption associated with time-restricted feeding did not impede motivation or acquisition of the task. In early PCA training (day 5), there were 2 goal-trackers (1 male, 1 female), 10 intermediates (5 male, 4 female), and 20 sign-trackers (10 male, 11 female). After 17 days there were no goal-trackers, 10 intermediates (4 male, 5 female), and 22 sign-trackers (12 male, 11 female). As expected with extended training, overall PCA scores shifted toward sign-tracking with extended training, indicating that the CS+ cue eventually acquires incentive salience even in rats that were initially goal-trackers or intermediates with negative PCA scores (Keefer et al., 2020; Villaruel & Chaudhri, 2016). Importantly, we also examined how PCA behavior is quantified and found many rats who do not exhibit strong lever pressing but instead display other lever-directed behavior sniffing, staring, biting, grabbing, or investigating the lever or cue light. This is particularly important for female rats as they display lower lever pressing overall compared to males. We applied the same analysis to food cup-directed behavior to examine behaviors beyond food cup head entry such as sniffing, staring, biting, grabbing, or investigating the food cup. Furthermore, we found total interaction time with either the food cup or CS+ cue and interaction bout length more indicative of respective behavioral phenotype than the number of interactions alone. We also assessed behavior during the inter-trial interval (ITI), and interestingly food cup-directed behavior during the ITI increased with extended training. Following extended PCA training, all rats were sensitive to satiety-induced devaluation regardless of sex, behavioral phenotype, or type of devaluation.

To better motivate learning, PCA is typically executed under daily food deprivation. However, rats often drop to 80-90% of their *ad libitum* body weight during this period (Bacharach et al., 2023; Bien & Smith, 2023; Derman & Lattal, 2023; Stapf et al., 2024). This raises concerns because PCA scores and behavioral phenotypes are greatly influenced by environmental factors, such as stress. Chronic food restriction, such as that used in typical PCA experiments, increases activation of the HPA axis, increases adrenal weight, and increases basal and stress-induced corticosterone levels (Kiss et al, 1994; Stewart et al, 1998; El Fazaa et al, 2000; Heiderstadt et al, 2000; Gursoy et al, 2001; Sobarenes-Chavez et al, 2024). These responses return to normal following a return to *ad libitum* access to food and are not observed when time-restricted feeding access to *ad libitum* food is used (Heiderstadt et al, 2000; Sobarenes-Chavez et al, 2024). Food restriction also increases stress-responsive behaviors such as anxiety-related behaviors, immobility in forced swim tasks, and conditioned fear (Pickens et al, 2010; Iio et al, 2015; Soberanes-Chavez et al, 2024). Interestingly, the food restriction-induced increases in anxiety-like behavior are not observed using time-restricted feeding regimens (Soberanes-Chavez et al, 2024). Importantly, acute and chronic food restriction is an effective stressor to increase drug-related behaviors such as cocaine-induced locomotor sensitization (Stamp et al, 2008; Allen et al, 2013) and heroin-seeking behavior (Shalev 2012; D’Cuhna et al, 2013; Sedki et al, 2013; Chisholm et al, 2020). In the context of PCA, stress has been shown to increase sign-tracking behavior (Lomanowska et al., 2011; Turfe et al., 2024; Spring et al, 2021) and therefore chronic food restriction could be impacting the overall distribution of PCA scores and behavioral phenotypes. Thus, we believed using a less severe time-restriction feeding paradigm that allowed rats to maintain weight eliminated food-restriction stress while still allowing rats to be in as fasted state daily prior to PCA training. Using our time-restricted feeding paradigm with a 4-hour daily feeding window, rats maintained *ad libitum* body weight and caloric intake and continued to increase in body weight throughout PCA training. Notably, even at *ad libitum* body weights, rats were unimpaired in learning the PCA task; within 3 days of training, rats successfully differentiated between the CS+ and CS-, a rate similar to rats trained under food deprivation (Bacharach et al., 2023; Bien & Smith, 2023; Derman & Lattal, 2023).

However, despite this less severe food restriction, most of the rats in the present experiment were sign-trackers by day 5 of training. A variety of factors influence PCA score so there may be factors beyond food deprivation stress that could be causing this bias towards sign-tracking behavior. Vendor and colony, for example, has been shown to play a role in PCA score (Fitzpatrick et al., 2013). Among Sprague-Dawley rats, another outbred rat strain commonly used in PCA, rats from Harlan were more likely to be sign-trackers and rats from Charles River were more likely to be goal-trackers (Fitzpatrick et al, 2013). Furthermore, in both Sprague-Dawley and Long-Evans rats, individual colonies within vendor also differed significantly in PCA behavior, indicating that both subtle genetic and environmental factors could influence incidence of sign-tracking and goal-tracking behavior (Fitzpatrick et al., 2013; Khoo et al., 2022). In the current experiment, only one cohort of animals was used from Envigo (formerly Harlan) and thus we cannot exclude the possibility that some of the distribution of PCA scores were impacted by our vendor selection and the individual colony that they originated from. Therefore, consideration of vendor and colony is an important variable to account for in the design of future PCA studies and these studies would benefit from the use of multiple cohorts to ensure the effects are not driven by an individual colony.

A majority of PCA studies quantify lever-directed and food-directed behavior through automatic recordings of lever presses and food-cup head entries (Bien & Smith, 2023; Derman & Matthew Lattal, 2023; Meyer et al., 2012; Stapf et al., 2024; Turfe et al., 2024). However, we observed behaviors during the cue presentation that may not trigger a lever press or head entry. This included sniffing, staring, grabbing, biting, and rearing. Notably, this behavior was directed not just at the food cup or lever, but also at the cue light. Including these behaviors in the analyses provided greater insight into the PCA phenotypes, particularly for the intermediate phenotype. Intermediate rats had a greater number of interactions with the lever and light than sign-trackers and interacted with the food cup as much as goal trackers though we did not include the goal trackers in the analysis due to the low n. However, it seems that intermediate rats had briefer bouts of interaction, switching frequently between the lever and food cup, whereas sign-trackers and goal-trackers often had long uninterrupted bouts of interaction with their preferred target. Indeed, interaction time seems to be best at delineating the three phenotypic groups compared to the number of interactions or the number of lever presses or food cup entries. There are differences in the time spent interacting with lever and time spent interacting with the food cup between all three phenotypes. However, it is important to note that goal-trackers had a small group size (n=2) and was therefore not included in the statistical analysis. Thus, there is a limit to the extent to which goal-tracking behavior can be reasonably interpreted from the present data. Nevertheless, these preliminary results, combined with the differences observed between sign-trackers and intermediates (both of which had more robust group sizes), suggest that our expanded behavior analysis improved identification of phenotypic groups during PCA.

Furthermore, we examined behavior during the inter-trial interval (ITI), a relatively unexplored period of PCA training. Rats interacted with the site of the CS+ lever and light, though there was no effect of phenotype (data not shown) or with duration of training. Time spent interacting with the food cup during the ITI also occurred and increased with extended training, particularly in female rats. It is likely that the association of the food cup with the sucrose reward and the conditioned context of the operant box were strengthened with extended training, which caused the increase in food cup time during the ITI (Fraser & Janak, 2019; Petrovich et al., 2007). Notably, this increased interaction with the food cup is restricted to the ITI; the increasing PCA score reflects greater lever/light interaction during CS+ presentation, suggesting the cue itself had even more incentive salience with extended PCA training (Flagel et al., 2009).

In animals with shorter duration PCA training, sign-trackers are resistant to both satiety-induced and aversion-induced reward devaluation (Keefer et al., 2020; Morrison et al., 2015). However, following extended training, sign-trackers no longer show resistance to satiety-induced reward devaluation (Bien & Smith, 2023; Keefer et al., 2020; Stapf et al., 2024). We have replicated these findings and have expanded upon these results to compare two different types of satiety-induced devaluation: satiety from chow and satiety from sucrose. Both forms of devaluation decreased CS+ responses to a similar degree. This change in behavioral flexibility may be due to changes in the neuronal mechanisms mediating PCA. For example, striatal dopamine release is observed in sign-trackers (but not goal-trackers) in response to cue presentation following acute training but declines with extended training; it is perhaps this activation that influences devaluation sensitivity (Keefer et al., 2020).

Varying effects of sex of have been observed on PCA acquisition, behavioral phenotype, and devaluation. Some studies report increased sign-tracking and lever pressing in female rats (Pitchers et al., 2015; Stapf et al., 2024), while others find no difference in acquisition or ultimate PCA score between males and females (Bien & Smith, 2023; Pitchers et al., 2015). The present study found no difference in distribution of PCA score with limited or extended training. Furthermore, we observed minimal sex differences in lever-directed behavior. However, we found that males had a greater amount of lever presses compared to females throughout the experiment. When behaviors beyond lever pressing (chewing, biting, staring, or otherwise investigating the light or lever) were included, sex differences were limited to the intermediate phenotype. Female intermediate rats interacted more with both the lever and food cup during cue presentation than male intermediate rats. No difference was observed between male and female sign-trackers or goal-trackers. However, during the inter-trial interval, female rats interacted with the food cup more than male rats suggesting greater goal-directed behavior in the absence of the reward-paired cue or greater context conditioning in female rats.

There are also discrepancies in whether there is an effect of sex on reward devaluation depending on the study. In one study there were no sex differences observed in reward devaluation (Bien & Smith, 2023), whereas others report that females were less sensitive to reward devaluation. In particular, one study found that female sign-trackers maintain insensitivity to reward devaluation following extended PCA training while male sign-trackers do not (Keefer et al., 2023; Stapf et al., 2024). However, following our extended training protocol, we saw no effect of sex on reward devaluation – both males and females showed similar sensitivity to satiety-induced devaluation though this may be due to strain differences as the prior studies used Sprague-Dawley rats and our study used Long-Evans rats. In support of this, the effect of sex on PCA behavior and devaluation vary across strain (Bien & Smith, 2023; Pitchers et al., 2015). For example, Sprague-Dawley rats show increased sign-tracking and devaluation resistance in females, while no sex difference is seen Heterogeneous Stock rats (Pitchers et al., 2015). However, there are conflicting reports on the role of sex in Long-Evans rats (Bien & Smith, 2023; Stapf et al., 2024). Furthermore, PCA behavior does not differ across estrous cycle (Pitchers et al., 2015). Taken together, it seems that the effect of sex on PCA behavior and devaluation, particularly in Long Evans rats, is minimal.

In the current study we have replicated previous findings that PCA scores trend towards sign-tracking with increased training and that sensitivity to reward devaluation increases with extended training. Furthermore, we contributed to the growing findings that sex differences in PCA behavior and reward devaluation seem to be minimal. It appears that any sex differences are concentrated in intermediate rats, a relatively unstudied phenotypic group. Although sign-trackers and goal-trackers are often the focus of Pavlovian studies, perhaps studying intermediate rats may provide further information about the encoding of incentive salience towards cues. We also expanded the types of behaviors that were used to establish PCA scores, examining previously unrecorded behaviors both during cue presentation and the inter-trial interval. Moreover, the data indirectly supports the influence of vendors and colony on PCA behavior. Future work should consider litter to avoid biasing PCA score toward sign-or goal-tracking. Taken together, the present work establishes a time restricted 4-hr feeding window to minimize weight loss during PCA training and defines a more robust way to analyze PCA phenotype by examining behaviors not typically studied.

## Data Statement

Data is available upon request

## Acknowledgements

This work was funded by the Department of Pharmacology, Physiology, and Neurobiology at the University of Cincinnati College of Medicine

## Author Contributions

ILR: Conceptualization, Formal Analysis, Investigation, Methodology, Writing; KB: Investigation; YMUL: Conceptualization, Formal Analysis, Methodology, Resources, Supervision, Writing – Original, Review, and Editing; JRM: Conceptualization, Formal Analysis, Methodology, Resources, Supervision, Writing – Original, Review, and Editing

## Supplemental Figures

**Figure S1.**
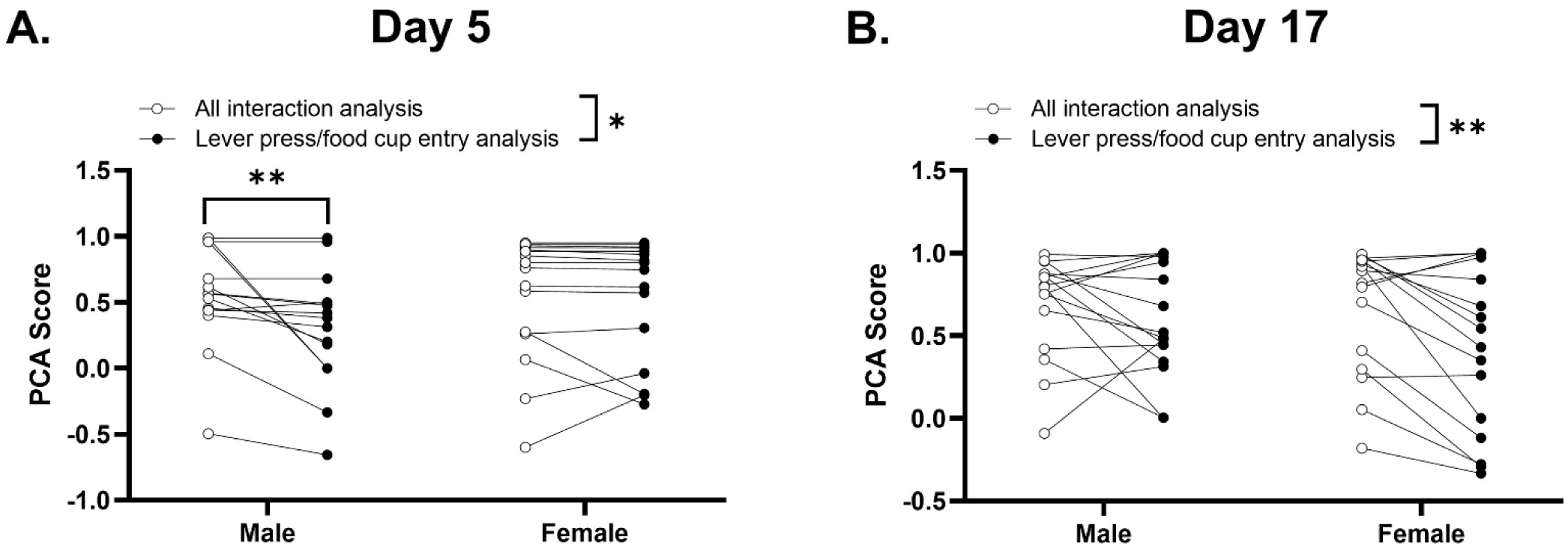
The overall distribution of PCA score was impacted by the method of analysis used to calculate the PCA score. A) PCA scores on day 5 of PCA training calculated by using the total time spent interacting with either the lever/light or the food cup that encompasses behavior beyond lever presses or head entry into the food cup (all interaction analysis) resulted in higher PCA scores than the standard approach using lever presses and food cup head entry. This effect was primarily observed in male, but not female, rats. This ultimately results in a greater number of sign tracker rats using the all interaction analysis approach. B) On day 17 of PCA training, the all interaction analysis approach to calculating PCA score resulted in significantly higher PCA scores in both male and female rats compared to using the lever press/food cup head entry analysis approach. The all interaction analysis approach ultimately results in a greater number of sign tracker rats compared to the lever press/food cup head entry analysis approach. Data are shown as mean ± SEM. *p<0.05, **p<0.01.

**Figure S2.**
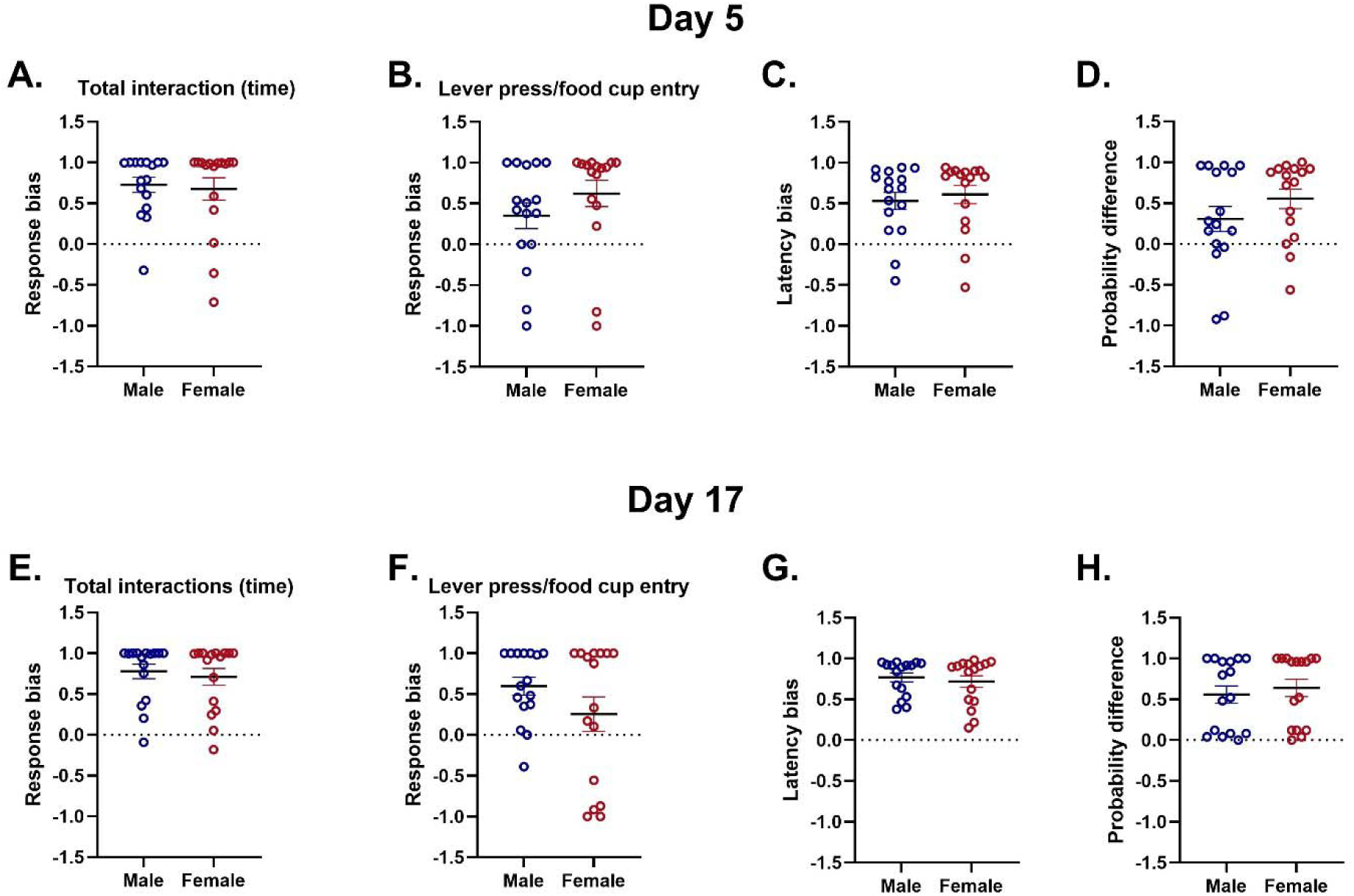
The individual metrics used to calculate PCA scores does not differ by sex in either early or late PCA training. PCA scores are calculated using response bias, latency bias, and probability difference in approach to the lever or food cup. A) The response bias calculated using the all interaction analysis (total time spent interacting with either the lever/light or the food cup) does not significantly differ between male and female rats in early PCA training (Day 5). B) The response bias calculated by using lever presses or food cup head entry does not significantly differ between male and female rats in early PCA training. C) Latency bias does not differ between male and female rats in early PCA training. D) Probability difference in approach to either the lever or food cup does not differ between male and female rats in early PCA training. E) Response bias calculated using the total interaction time with either the lever/light or food cup does not differ between male and female rats in late PCA training (Day 17). F) Response bias calculated using lever presses or food cup head entry does not differ between male and female rats in late PCA training. G) Latency bias does not differ between male and female rats in late PCA training. H) Probability difference in approach to either the lever or food cup does not differ between male and female rats in late PCA training. Data are shown as mean ± SEM.

**Figure S3.**
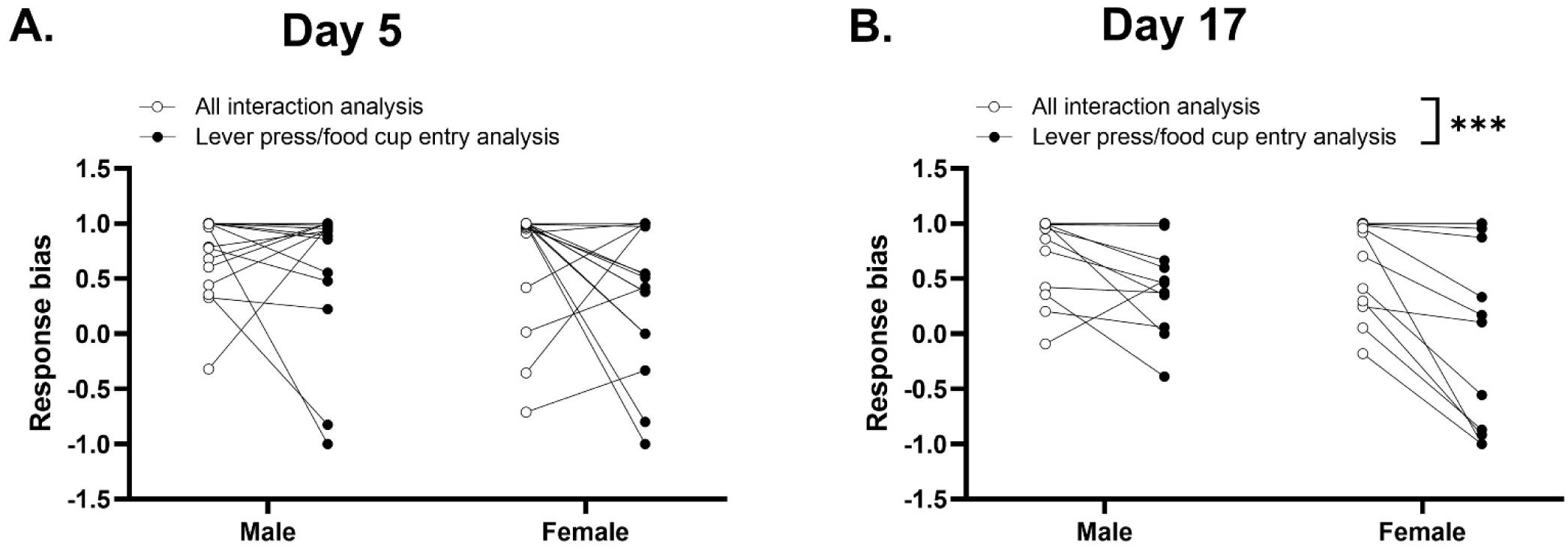
The calculation of response balance is impacted by the method of analysis used in late PCA training. A) There is no significant difference in the response bias value as calculated using the all interaction analysis approach compared to the lever press/food cup entry analysis approach on day 5 of PCA training in either male or female rats. B) Response bias calculated using the all interaction analysis approach is significantly higher compared to using the lever press/food cup entry analysis approach in male and female rats on day 17 of PCA training. Data are shown as mean ± SEM. ***p<0.001.

## References

1. Allen C. P., Zhou Y., Leri F. (2013). Effect of food restriction on cocaine locomotor sensitization in Sprague-Dawley rats: role of kappa opioid receptors. Psychopharmacology (Berl), 226(3): 571–578. DOI: 10.1007/s00213-012-2930-7.

2. Bacharach, S. Z., Martin, D. A., Stapf, C. A., Sun, F., Li, Y., Cheer, J. F., & Calu, D. J. (2023). Decreased Ventral Tegmental Area CB1R Signaling Reduces Sign Tracking and Shifts Cue– Outcome Dynamics in Rat Nucleus Accumbens. The Journal of Neuroscience, 43(25), 4684– 4696. 10.1523/jneurosci.1486-22.2023

3. Bien, E., & Smith, K. (2023). The role of sex on sign-tracking acquisition and outcome devaluation sensitivity in Long Evans rats. Behavioural Brain Research, 455(July), 114656. 10.1016/j.bbr.2023.114656

4. Chisholm A., Iannuzzi J., Rizzo D., Gonzalez N., Fortin E., Bumbu A., Batallan Burrowes A. A., Chapman C. A., Shalev U. (2020). The role of the paraventricular nucleus of the thalamus in the augmentation of heroin seeking induced by chronic food restriction. Addiction Biology, 25(2): e12708. DOI: 10.1111/adb.12708.

5. D’Cunha T. M., Sedki F., Macri J., Casola C., Shalev U. (2013). The effects of chronic food restriction on cue-induced heroin seeking in abstinent male rats. Psychopharmacology (Berl), 225(1): 241–250. DOI: 10.1007/s00213-012-2810-1.

6. Derman, R. C., & Matthew Lattal, K. (2023). Acute stress persistently alters instrumental motivation without affecting appetitive Pavlovian conditioning, extinction, or contextual renewal. Neurobiology of Learning and Memory, 202(January), 107771. 10.1016/j.nlm.2023.107771

7. El Fazaa S., Gharbi N., Kamoun A., Somody L. (2000). Vasopressin and A1 noradrenaline turnover during food or water deprivation in the rat. Comp Biochem Physiol C Toxicol Pharmacol, 126(2): 129–137. DOI: 10.1016/s0742-8413(00)00105-5.

8. Fitzpatrick, C. J., Gopalakrishnan, S., Cogan, E. S., Yager, L. M., Meyer, P. J., Lovic, V., Saunders, B. T., Parker, C. C., Gonzales, N. M., Aryee, E., Flagel, S. B., Palmer, A. A., Robinson, T. E., & Morrow, J. D. (2013). Variation in the Form of Pavlovian Conditioned Approach Behavior among Outbred Male Sprague-Dawley Rats from Different Vendors and Colonies: Sign-Tracking vs. Goal-Tracking. PLoS ONE, 8(10). 10.1371/journal.pone.0075042

9. Flagel, S. B., Akil, H., & Robinson, T. E. (2009). Individual differences in the attribution of incentive salience to reward-related cues: Implications for addiction. Neuropharmacology, 56(SUPPL. 1), 139–148. 10.1016/j.neuropharm.2008.06.027

10. Fraser, K. M., & Janak, P. H. (2019). Occasion setters attain incentive motivational value: Implications for contextual influences on reward-seeking. Learning and Memory, 26(8), 291–298. 10.1101/lm.049320.119

11. Friard, O., Gamba, M., (2016). BORIS : a free, versatile open-source event-logging software for video/audio coding and live observations. Methods in Ecology and Evolution, 7, 1325–1330.. 10.1111/2041-210x.12584

12. Gursoy E., Cardounel A., Hu Y., Kalimi M. (2001). Biological effects of long-term caloric restriction: adaptation with simultaneous administration of caloric stress plus repeated immobilization stress in rats. Exp Biol Med, 226(2): 97–102. DOI: 10.1177/153537020122600206.

13. Heiderstadt K. M., McLaughlin R. M., Wright D. C., Walker S. E., Gomez-Sanchez C. E. (2000). The effect of chronic food and water restriction on open-field behaviour and serum corticosterone levels in rats. Lab Anim, 34(1): 20–28. DOI: 10.1258/002367700780578028.

14. Horstmann, A., Dietrich, A., Mathar, D., Pössel, M., Villringer, A., & Neumann, J. (2015). Slave to habit? Obesity is associated with decreased behavioural sensitivity to reward devaluation. Appetite, 87, 175–183. 10.1016/j.appet.2014.12.212

15. Hou S., Glover E. J. (2022). Pi USB Cam: A Simple and Affordable DIY Solution That Enables High-Quality, High-Throughput Video Capture for Behavioral Neuroscience Research. eNeuro, 9(5): DOI: 10.1523/ENEURO.0224-22.2022. PMC9522465

16. Iio W., Tokutake Y., Koike H., Matsukawa N., Tsukahara T., Chohnan S., Toyoda A. (2015). Effects of chronic mild food restriction on behavior and the hypothalamic malonyl-CoA signaling pathway. Anim Sci J, 86(2): 181–188. DOI: 10.1111/asj.12255.

17. Keefer, S. E., Bacharach, S. Z., Kochli, D. E., Chabot, J. M., & Calu, D. J. (2020). Effects of Limited and Extended Pavlovian Training on Devaluation Sensitivity of Sign- and Goal-Tracking Rats. Frontiers in Behavioral Neuroscience, 14(February), 1–13. 10.3389/fnbeh.2020.00003

18. Keefer, S. E., Richard, J. M., & Richard, J. M. (2023). Sex-biased effects of outcome devaluation by sensory-specific satiety on Pavlovian-conditioned behavior. October, 1–13. 10.3389/fnbeh.2023.1259003

19. Khoo, S. Y.-S., Uhrig, A., Samaha, A.-N., & Chaudhri, N. (2022). Does vendor breeding colony influence sign- and goal-tracking in Pavlovian conditioned approach? A preregistered empirical replication. Neuroanatomy and Behaviour, 4, e46–e46. 10.35430/nab.2022.e46

20. Kiss A., Jezova D., Aguilera G. (1994). Activity of the hypothalamic pituitary adrenal axis and sympathoadrenal system during food and water deprivation in the rat. Brain Res 663(1): 84–92. DOI: 10.1016/0006-8993(94)90465-0.

21. Kuhn, B., Campus, P., & Flagel, S. (2018). The Neurobiological Mechanisms Underlying Sign-Tracking Behavior. *Sign-Tracking and Drug Addiction*, November, 35–74.

22. Lomanowska, A. M., Lovic, V., Rankine, M. J., Mooney, S. J., Robinson, T. E., & Kraemer, G. W. (2011). Inadequate early social experience increases the incentive salience of reward-related cues in adulthood. Behavioural Brain Research, 220(1),

23. Marzuki, A. A., Banca, P., Garofalo, S., Degni, L. A. E., Dalbagno, D., Badioli, M., Sule, A., Kaser, M., Conway-Morris, A., Sahakian, B. J., & Robbins, T. W. (2024). Compulsive avoidance in youths and adults with OCD: an aversive pavlovian-to-instrumental transfer study. Translational Psychiatry, 14(1), 1–12. 10.1038/s41398-024-03028-1

24. Meyer, P. J., Lovic, V., Saunders, B. T., Yager, L. M., Flagel, S. B., Morrow, J. D., & Robinson, T. E. (2012). Quantifying individual variation in the propensity to attribute incentive salience to reward cues. PLoS ONE, 7(6). 10.1371/journal.pone.0038987

25. Morrison, S. E., Bamkole, M. A., & Nicola, S. M. (2015). Sign tracking, but not goal tracking, is resistant to outcome devaluation. Frontiers in Neuroscience, 9(DEC), 1–12. 10.3389/fnins.2015.00468

26. Petrovich, G. D., Ross, C. A., Gallagher, M., & Holland, P. C. (2007). Learned contextual cue potentiates eating in rats. Physiology and Behavior, 90(2–3), 362–367. 10.1016/j.physbeh.2006.09.031

27. Pickens C. L., Navarre B. M., Nair S. G. (2010). Incubation of conditioned fear in the conditioned suppression model in rats: role of food-restriction conditions, length of conditioned stimulus, and generality to conditioned freezing. Neuroscience, 169(4): 1501–1510. DOI: 10.1016/j.neuroscience.2010.06.036. PMC2920731

28. Pitchers, K. K., Flagel, S. B., O’Donnell, E. G., Solberg Woods, L. C., Sarter, M., & Robinson, T. E. (2015). Individual variation in the propensity to attribute incentive salience to a food cue: Influence of sex. Behavioural Brain Research, 278, 462–469. 10.1016/j.bbr.2014.10.036

29. Sedki F., Abbas Z., Angelis S., Martin J., D’Cunha T., Shalev U. (2013). Is it stress? The role of stress related systems in chronic food restriction-induced augmentation of heroin seeking in the rat. Front Neurosci 7: 98. DOI: 10.3389/fnins.2013.00098. PMC3674335

30. Shalev U. (2012). Chronic food restriction augments the reinstatement of extinguished heroin-seeking behavior in rats. Addiction Biology 17(4): 691–693. DOI: 10.1111/j.1369-1600.2010.00303.x.

31. Soberanes-Chavez P., Trujillo-Barrera J., de Gortari P. (2024). Circadian Synchronization of Feeding Attenuates Rats’ Food Restriction-Induced Anxiety and Amygdalar Thyrotropin-Releasing Hormone Downregulation. Int J Mol Sci, 25(11): DOI: 10.3390/ijms25115857. PMC11172148

32. Spring, M. G., Caccamise, A., Panther, E. A., Windsor, B. M., Soni, K. R., McReynolds, J. R., Wheeler, D. S., Mantsch, J. R., & Wheeler, R. A. (2021). Chronic stress prevents cortico-accumbens cue encoding and alters conditioned approach. Journal of Neuroscience, 41(11), 2428–2436. 10.1523/JNEUROSCI.1869-20.2021

33. Stamp J. A., Mashoodh R., van Kampen J. M., Robertson H. A. (2008). Food restriction enhances peak corticosterone levels, cocaine-induced locomotor activity, and DeltaFosB expression in the nucleus accumbens of the rat. Brain Research, 1204(94–101. DOI: 10.1016/j.brainres.2008.02.019.

34. Stapf, C. A., Keefer, S. E., McInerney, J. M., & Calu, D. J. (2024). Sex specific effects of dorsomedial striatal cannabinoid receptor-1 signaling on Pavlovian outcome devaluation. BioRxiv, 2024.05.01.592059. 10.1101/2024.05.01.592059

35. Stewart J., Meaney M. J., Aitken D., Jensen L., Kalant N. (1988). The effects of acute and life-long food restriction on basal and stress-induced serum corticosterone levels in young and aged rats. Endocrinology, 123(4): 1934–1941. DOI: 10.1210/endo-123-4-1934.

36. Swintosky, M., Brennan, J. T., Koziel, C., Paulus, J. P., & Sara, E. (2022). Sign tracking predicts suboptimal behavior in a rodent gambling task. 238(9), 2645–2660.

37. Turfe, A., Westbrook, S. R., Lopez, S. A., Chang, S. E., & Flagel, S. B. (2024). The effect of corticosterone on the acquisition of Pavlovian conditioned approach behavior in rats is dependent on sex and vendor. Hormones and Behavior, 164(March), 105609. 10.1016/j.yhbeh.2024.105609

38. Villaruel, F. R., & Chaudhri, N. (2016). Individual differences in the attribution of incentive salience to a pavlovian alcohol cue. Frontiers in Behavioral Neuroscience, 10(DEC), 1–13. 10.3389/fnbeh.2016.00238

